# A protocol for an intercomparison of biodiversity and ecosystem services models using harmonized land-use and climate scenarios

**DOI:** 10.1101/300632

**Authors:** HyeJin Kim, Isabel M.D. Rosa, Rob Alkemade, Paul Leadley, George Hurtt, Alexander Popp, Detlef P van Vuuren, Peter Anthoni, Almut Arneth, Daniele Baisero, Emma Caton, Rebecca Chaplin-Kramer, Louise Chini, Adriana De Palma, Fulvio Di Fulvio, Moreno Di Marco, Felipe Espinoza, Simon Ferrier, Shinichiro Fujimori, Ricardo E. Gonzalez, Maya Gueguen, Carlos Guerra, Mike Harfoot, Thomas D. Harwood, Tomoko Hasegawa, Vanessa Haverd, Petr Havlík, Stefanie Hellweg, Samantha L. L. Hill, Akiko Hirata, Andrew J. Hoskins, Jan H. Janse, Walter Jetz, Justin A. Johnson, Andreas Krause, David Leclère, Ines S. Martins, Tetsuya Matsui, Cory Merow, Michael Obersteiner, Haruka Ohashi, Benjamin Poulter, Andy Purvis, Benjamin Quesada, Carlo Rondinini, Aafke Schipper, Richard Sharp, Kiyoshi Takahashi, Wilfried Thuiller, Nicolas Titeux, Piero Visconti, Christopher Ware, Florian Wolf, Henrique M. Pereira

## Abstract

To support the assessments of the Intergovernmental Science-Policy Platform on Biodiversity and Ecosystem Services (IPBES), the IPBES Expert Group on Scenarios and Models is carrying out an intercomparison of biodiversity and ecosystem services models using harmonized scenarios (BES-SIM). The goals of BES-SIM are (1) to project the global impacts of land use and climate change on biodiversity and ecosystem services (i.e. nature’s contributions to people) over the coming decades, compared to the 20^th^ century, using a set of common metrics at multiple scales, and (2) to identify model uncertainties and research gaps through the comparisons of projected biodiversity and ecosystem services across models. BES-SIM uses three scenarios combining specific Shared Socio-economic Pathways (SSPs) and Representative Concentration Pathways (RCPs) to explore a wide range of land-use change and climate change futures. This paper describes the rationale for scenarios selection, the process of harmonizing input data for land use, based on the second phase of the Land Use Harmonization Project (LUH2), and climate, the biodiversity and ecosystem service models used, the core simulations carried out, the harmonization of the model output metrics, and the treatment of uncertainty. The results of this collaborative modelling project will support the ongoing global assessment of IPBES, strengthen ties between IPBES and the Intergovernmental Panel on Climate Change (IPCC) scenarios and modelling processes, advise the Convention on Biological Diversity (CBD) on its development of a post-2020 strategic plans and conservation goals, and inform the development of a new generation of nature-centred scenarios.

## 1 Introduction

Understanding how anthropogenic activities impact biodiversity, ecosystems, and human societies is essential for nature conservation and sustainable development. Land use and climate change are widely recognized as two of the main drivers of future biodiversity change (Hirsch and CBD, 2010; Maxwell et al., 2016; Sala, 2000; CBD and UNEP, 2014) with potentially severe impacts on ecosystem services and ultimately human well-being (Cardinale et al., 2012; MA, 2005). Habitat and land-use changes, resulting from past, present and future human activities, have immediate impacts on biodiversity and ecosystem services whereas the impacts of climate change have considerable lag times (Lehsten et al., 2015). Therefore, current and future land-use projections are essential elements for assessing biodiversity and ecosystem change (Titeux et al., 2016, 2017). Climate change has already observed to have direct and indirect impact on biodiversity and ecosystems and it is projected to intensify as we approach the end of the century with potentially severe consequences on species and habitats, thereby also on ecosystem functions and ecosystem services at high levels of climate change (Pecl et al., 2017; Settele et al., 2015).

Global environmental assessments, such as the Millennium Ecosystem Assessment (MA 2005), the Global Biodiversity Outlooks (GBO), the multiple iterations of the Global Environmental Outlook (GEO), the Intergovernmental Panel on Climate Change (IPCC), and other studies have used scenarios to assess the impact of socio-economic development pathways on land use and climate and their consequences for biodiversity and ecosystem services (Jantz et al., 2015; Pereira et al., 2010). Models are used in quantifying the narratives of scenarios using selected and modellable drivers, which describe key components of a system or relationships between them (Ferrier et al. 2016). So far, these scenarios analysis exercises have been based on a single model or a small number of models, and cross-model harmonization and uncertainty analysis have been limited. The Expert Group on Scenarios and Models of the Intergovernmental Science-Policy Platform on Biodiversity and Ecosystem Services (IPBES) is addressing this issue by carrying out a biodiversity and ecosystem services model intercomparison with harmonized scenarios.

Over the last two decades, IPCC has fostered the development of global scenarios to inform climate mitigation and adaptation policies. The Representative Concentration Pathways (RCPs) describe different climate futures based on greenhouse gas emissions over the 21^st^ century (van Vuuren et al., 2011). These emissions pathways have been converted into climate projections in the most recent Climate Model Inter-comparison Project (CMIP5). In parallel, the climate research community also developed the Shared Socio-economic Pathways (SSPs) which consist of trajectories of future human development with different socio-economic conditions and associated land-use projections (Popp et al., 2017; Riahi et al., 2017). The SSPs can be combined with RCP-based climate projections to explore a range of futures for climate change and land-use change and are being used in a wide range of impact modelling intercomparisons (Rosenzweig et al., 2017; van Vuuren et al., 2014). Therefore, the use of the SSP-RCP framework for modelling the impacts on biodiversity and ecosystem services provides an outstanding opportunity to build bridges between the climate, biodiversity and ecosystem services communities, and has been explicitly recommended as a research priority in the IPBES assessment on scenarios and models (Ferrier et al. 2016).

Model intercomparisons bring together different communities of practice for comparable and complementary modelling, in order to improve the robustness and comprehensiveness of the subject modelled, and to estimate associated uncertainties (Warszawski et al., 2014). In the last decades, various model intercomparison projects (MIPs) have been initiated to assess the magnitude and uncertainty of climate change impacts. For instance, the Inter-Sectoral Impact Model Intercomparison Project (ISI-MIP) was initiated in 2012 to quantify and synthesize climate change impacts across sectors and scales (Frieler et al., 2015; Rosenzweig et al., 2017). The ISI-MIP aims to bridge sectors such as agriculture, forestry, fisheries, water, energy, and health with Global Circulation Models (GCMs), Earth System Models (ESMs), and Integrated Assessment Models (IAMs) for more integrated and impact-driven modelling and assessment (Frieler et al., 2017).

Here, we present the methodology used to carry out a Biodiversity and Ecosystem Services Scenario-based Intercomparison of Models (BES-SIM) in terrestrial and freshwater ecosystems. The BES-SIM project addresses the following questions: (1) What are the projected magnitudes and spatial distribution of biodiversity and ecosystem services under a range of climate and land-use future scenarios? (2) What is the magnitude of the uncertainties associated with the projections obtained from different models and scenarios? We brought together ten biodiversity models and six ecosystem functions and ecosystem services models to assess impacts of land-use and climate change scenarios in coming decades (up to 2070) and to hindcast changes to the last century (to 1900). The modelling approaches differ in several ways in how they treat biodiversity and ecosystem services responses to land use and climate changes, including the use of correlative, deductive, and process-based approaches, and in how they treat spatial scale and temporal dynamics. We assess different dimensions of biodiversity including species richness, species abundance, community composition, and habitat shifts, as well as a range of measures on ecosystem services such as food production, pollination, water quantity and quality, climate regulation, soil protection, and pest control. This paper provides an overview of the scenarios, models and metrics used in this intercomparison, thus a roadmap for further analyses that is envisaged to be integrated into the first global assessment of the IPBES (Figure 1).

**Figure 1:**
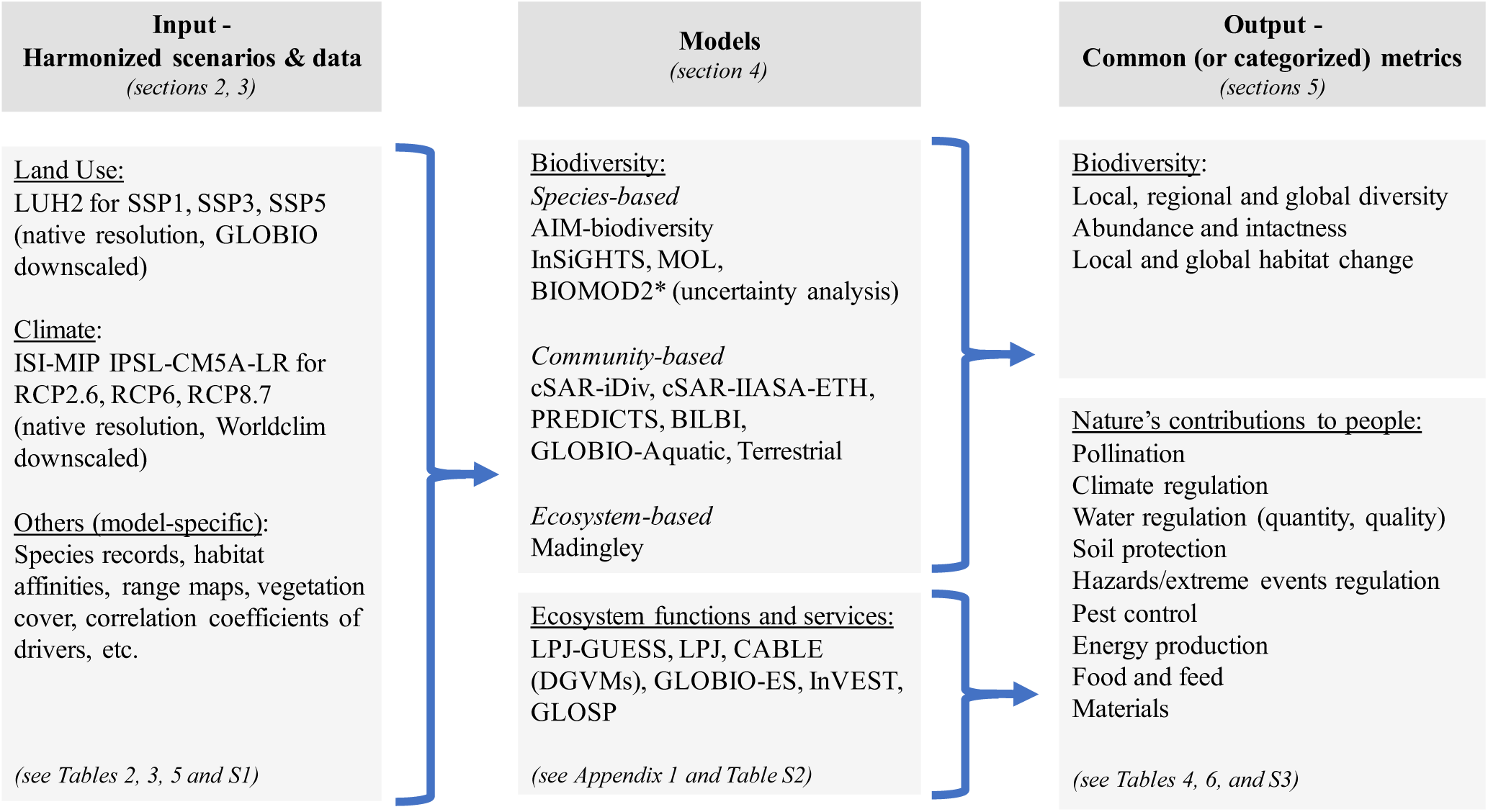
Input-models-output flowchart of BES-SIM.

## 2 Scenarios selection

All the models involved in BES-SIM used the same set of scenarios using particular combinations of SSPs and RCPs. In the selection of the scenarios, we used the following criteria: 1) data on projections should be readily available, and 2) the total set should cover a broad range of land-use change and climate change projections. The first criterion implied that we selected SSP-RCP combinations that are included in the ScenarioMIP protocol as part of CMIP6 (O’Neill et al., 2016), as harmonised data was available for these runs and these form the basis of the CMIP climate simulations. The second criteria implied a selection within the ScenarioMIP set of scenarios with low and high degrees of climate change and different land-use scenarios. Our final selection was SSP1 with RCP2.6 (moderate land-use pressure and low level of climate change) (van Vuuren et al., 2017), SSP3 with RCP6.0 (high land-use pressure and moderately high level of climate change) (Fujimori et al., 2017), and SSP5 with RCP8.5 (medium land-use pressure and very high level of climate change) (Kriegler et al., 2017), thus allowing us to assess a broad range of plausible futures (Table 1). Further, by combining projections of low and high anthropogenic pressure of land-use with low and high level of climate change projections, we can test these drivers’ individual and synergistic impacts on biodiversity and ecosystem services.

**Table 1:** Characteristics of (a) SSP and (b) RCP scenarios simulated in BES-SIM. (adopted from Moss et al., 2010; O’Neill et al., 2017; Popp et al., 2017; van Vuuren et al., 2011)

|  | SSP1 Sustainability | SSP3 Regional Rivalry | SSP5 Fossil-fuel Development |
| --- | --- | --- | --- |
| Population growth | Relatively low | Low (OECD countries) to high (high fertility countries) | Relatively low |
| Urbanization | High | Low | High |
| Equity and social cohesion | High | Low | High |
| Economic growth | High to medium | Slow | High |
| International trade and globalization | Moderate | Strongly constrained | High |
| Land-use regulation | Strong to avoid environmental trade-off | Low with continued deforestation due to agriculture expansion | Medium with slow decline in deforestation |
| Agricultural productivity | High improvements with diffusion of best practices | Low with slow technology development and restricted trade | Highly managed and resource intensive |
| Consumption & diet | Low growth in consumption, low-meat | Resource-intensive consumption | Material-intensive consumption, meat-rich diet |
| Environment | Improving | Serious degradation | Highly successful management |
| Carbon intensity | Low | High | High |
| Energy intensity | Low | High | High |
| Technology development | Rapid | Slow | Rapid |
| Institution effectiveness | Effective | Weak | Increasingly effective |
| Policy focus | Sustainable development | Security | Development, free market, human capital |
| Participation of the land-use sector in mitigation policies | Full | Limited | Full |
| International cooperation for climate change mitigation | No delay | Heavy delay | Delay |

(b) RCP scenarios
|  | RCP2.6 Low emissions | RCP6.0 Intermediate emissions | RCP8.5 High emissions |
| --- | --- | --- | --- |
| Radiative forcing | Peak at 3W/m<sup>2</sup> before 2100 and decline | Stabilizes without overshoot pathways to 6W/m<sup>2</sup> in 2100 | Rising forcing pathways leading to 8.5 W/m<sup>2</sup> in 2100 |
| Concentration (p.p.m) | Peak at 490 CO<sub>2</sub> equiv. before 2100 and then declines | 850 CO<sub>2</sub> equiv. (at stabilization after 2100) | >1,370 CO<sub>2</sub> equiv. in 2100 |
| Methane emission | Reduced | Stable | Rapid increase |
| Reliance on fossil fuels | Decline | Heavy | Heavy |
| Energy intensity | Low | Intermediate | High |
| Climate policies | Stringent |  | No implementation |

The first scenario (SSP1xRCP2.6) is characterized by relatively “environmentally-friendly world” with a low population growth, a relatively low demand for animal products, a high urbanization rate and a high agricultural productivity. These factors together lead to a decrease in land use of around 700 Mha globally over time (mostly pastures). This scenario is also characterised by low air pollution, while policies are introduced to limit the increase of greenhouse gases in the atmosphere, leading to an additional forcing of 2.6 W/m^2^ before 2100. The second scenario (SSP3xRCP6.0) is characterised by “regional rivalry”, leading high population growth, slow economic development, material-intensive consumption and low food demand per capita. Agricultural land intensification is low, especially due to very limited transfer of new agricultural technologies to developing countries. This scenario has land-use change hardly regulated, with large conversion of land to human-dominated uses, and has a relatively high level of climate change with radiative forcing of 6.0 W/m^2^ by 2100. The third scenario (SSP5xRCP8.5) is a world characterised by “strong economic growth” fuelled by fossil fuels, with low population growth, a high food demand per capita, a high urbanization rate but also a high agricultural productivity. As a result, there is a modest increase in land use. Air pollution policies are stringent, motivated by local health concerns. This scenario leads to a very high level of climate change with a radiative forcing of 8.5 W/m^2^ by 2100. Full descriptions of each SSP scenario are given in Popp et al. (2017) and Riahi et al. (2017).

## 3 Input data

A consistent set of land use and climate data was used across the models to the extent possible, using existing datasets. All models in BES-SIM used the newly released Land Use Harmonization dataset version 2 (LUH2, Hurtt et al., 2018). For the models that require climate data, we selected the climate projections of the past, present and future from CMIP5 / ISI-MIP2a (McSweeney and Jones, 2016) and its downscaled version from the WorldClim (Fick and Hijmans, 2017), as well as MAGICC 6.0 (Meinshausen et al., 2011a, 2011b) from the IMAGE model for GLOBIO models (Table 2). A complete list of input datasets and variables used by the models is documented in Table S1 of the Supplementary Materials.

**Table 2:** Sources of input data in BES-SIM.

| BES-SIM model | Land-use data<br>LUH2 v2.0 |  | Climate data |  |  |
| --- | --- | --- | --- | --- | --- |
|  | Native resolution<br>*0.25 degree* | Downscaled<br>(GLOBIO)<br>*300m* | CMIP5-IPSL<br>Native<br>Resolution<br>*0.5 degree* | CMIP5-IPSL<br>Downscaled<br>(WorldClim)<br>*1km* | IMAGE†<br>(MAGICC 6.0) |
| **Species-based models of biodiversity** |  |  |  |  |  |
| AIM-biodiversity | * |  | * |  |  |
| InSiGHTS | * |  |  | * |  |
| MOL | * |  |  | * |  |
| **Community-based models of biodiversity** |  |  |  |  |  |
| cSAR-iDiv | * |  |  |  |  |
| cSAR-IIASA-ETH | * |  |  |  |  |
| BILBI | * |  |  | * |  |
| PREDICTS | * |  |  |  |  |
| GLOBIO - Aquatic | * |  |  |  | * |
| GLOBIO4 - Terrestrial |  | * |  |  | * |
| **Ecosystems-based model of biodiversity** |  |  |  |  |  |
| Madingley | * |  | * |  |  |
| **Models of ecosystem functions and services** |  |  |  |  |  |
| LPJ-GUESS | * |  | * |  |  |
| LPJ | * |  | * |  |  |
| CABLE | * |  | * |  |  |
| GLOBIO-ES | * |  |  |  | * |
| InVEST |  | * |  | * |  |
| GLOSP | * |  | * |  |  |

†All GLOBIO models use MAGICC climate data from the IMAGE model.

### 3.1 Land cover and land-use change data

The land-use scenarios provide an assessment of land-use dynamics in response to a range of socio-economic drivers and their consequences for the land system. The IAMs used to model land-use scenarios – IMAGE for SSP1/RCP2.5, AIM for SSP3/RCP7.0, and REMIND/MAgPIE for SSP5/RCP8.0 – include different economic and land-use modules for the translation of narratives into consistent quantitative projections across scenarios (Popp et al., 2017). It is important to note that the land-use scenarios used, although driven mostly by the SSP storylines, were projected to be consistent with the paired RCPs and include biofuel deployment to mitigate climate change. As there was no land-use projection for SSP3 with RCP6.0, we chose the available closest simulation SSP3/RCP7.0 from the LUH2 datasets

The land-use projections from each of the IAMs was harmonized using the LUH2 methodology. LUH2 was developed for CMIP6 and provides a global gridded land-use dataset comprising estimates of historical land-use change (850-2015) and future projections (2015-2100), obtained by integrating and harmonizing land-use history with future projections of different IAMs (Jungclaus et al., 2017; Lawrence et al., 2016; O’Neill et al., 2016). Compared to the first version of the LUH (Hurtt et al., 2011), LUH2 (Hurtt et al., 2018) is driven by the latest SSPs, has a higher spatial resolution (0.25 vs 0.50 degree), more detailed land-use transitions (12 versus 5 possible land-use states), and increased data-driven constraints (Heinimann et al., 2017; Monfreda et al., 2008). LUH2 provides over 100 possible transitions per grid cell per year (e.g., crop rotations, shifting cultivation, agricultural changes, wood harvest) and various agricultural management layers (e.g., irrigation, synthetic nitrogen fertilizer, biofuel crops), all with annual time steps. The 12 states of land include the separation of primary and secondary natural vegetation into forest and non-forest sub-types, pasture into managed pasture and rangeland, and cropland into multiple crop functional types (C3 annual, C3 perennial, C4 annual, C4 perennial, and N fixing crops) (Table 3).

**Table 3:** Improvements made in the Land Use Harmonization v2 (LUH2) dataset from LUH v1. (Hurtt et al., 2011)

|  | LUH v1 | LUH v2 |
| --- | --- | --- |
| Spatial resolution | 0.5 degree | 0.25 degree |
| Time steps | Annually from 1500 to 2100 | Annually from 850 to 2100 |
| Land use categories | 5 categories<br>Primary<br>Secondary<br>Pasture<br>Urban<br>Crop | 12 categories<br>Forested primary land (primf)<br>Non-forested primary land (primn)<br>Potentially forested secondary land (secdf)<br>Potentially non-forested secondary land (secdn)<br>Managed pasture (pastr)<br>Rangeland (range)<br>Urban land (urban)<br>C3 annual crops (c3ann)<br>C3 perennial crops (c3per)<br>C4 annual crops (c4ann)<br>C4 perennial crops (c4per)<br>C3 nitrogen-fixing crops (c3nfx) |
| Future | RCPs (4)<br>2.6<br>4.5<br>6.0<br>8.5 | SSPs (6)<br>SSP1-RCP2.6<br>SSP4-RCP3.4<br>SSP2-RCP4.5<br>SSP4-RCP6.0<br>SSP3-RCP7.0<br>SSP5-RCP8.5 |
| Land use transitions | <20 per grid cell per year | >100 per grid cell per year |
| Improvements |  | - New shifting cultivation algorithm<br>- Landsat F/NF change constraint<br>- Expanded diagnostic package<br>- New historical wood harvest reconstruction<br>- Agricultural management layers: irrigation, fertilizer, biofuel crops, wood harvest product split, crop rotations, flooded (rice) |

For biodiversity and ecosystem services models that rely on discrete, high-resolution land-use data (i.e., the GLOBIO model for terrestrial biodiversity and the InVEST model), the fractional LUH2 data were downscaled to discrete land-use grids (10 arc-seconds resolution; ~300 m) with the land-use allocation routine of the GLOBIO4 model. To that end, the areas of urban, cropland, pasture, rangeland and forestry from LUH2 were first aggregated across the LUH2 grid cells to the regional level of the IMAGE model, with forestry consisting of the wood harvest from forested cells and non-forested cells with primary vegetation. Next, the totals per region were allocated to 300m cells with the GLOBIO4 land allocation routine, with specific suitability layers for urban, cropland, pasture, rangeland, and forestry. After allocation, cropland was reclassified into three intensity classes (low, medium, high) based on the amount of fertilizer per grid cell. More details on the downscaling procedure are provided in Annex 1 of the Supplement Materials.

### 3.2 Climate data

General Circulation Models (GCMs) are based on fundamental physical processes (e.g., conservation of energy, mass, and momentum and their interaction with the climate system) and simulate climate patterns of temperature, precipitation and extreme events at a large scale (Frischknecht et al., 2016). Some GCMs now incorporate elements of Earth’s climate system (e.g. atmospheric chemistry, soil and vegetation, land and sea ice, carbon cycle) in ESMs (GCM with interactive carbon cycle), and have dynamically downscaled models with higher resolution data in Regional Climate Models (RCMs).

A large number of climate datasets are available today from multiple GCMs, but not all GCMs provide projections for all RCPs. Moreover, some models in BES-SIM require continuous time-series data. In order to harmonize the climate date to be used across biodiversity and ecosystem service models, we chose the bias-corrected climate projections from CMIP5, which were also adopted by ISIMIP2a (Hempel et al., 2013) or their downscaled versions available from the WorldClim (Fick and Hijmans, 2017). Most analysis were carried out using a single GCM, the IPSL-CM5A-LR (Dufresne et al., 2013), to avoid a random selection of GCMs by the different teams (Table 2).

The ISI-MIP fast-track output from the IPSL model provides 12 climate variables on daily time steps from pre-industrial period 1951 to 2099 at 0.5-degree resolution (McSweeney and Jones, 2016). The WorldClim downscaled dataset has 19 bioclimatic variables derived from monthly temperature and rainfall for 1960-1990 with multi-year averages for specific points in time (e.g., 2050, 2070) up to 2070. Six models in BES-SIM used ISI-MIP2a dataset and three models used WorldClim. An exception was made to the GLOBIO models, which used MAGICC 6.0 climate data (Meinshausen et al., 2011b, 2011a) in the IMAGE model framework (Stehfest et al., 2014), to which GLOBIO is tightly connected (Table 2). The variables used from climate dataset in each model are listed in Table S1.

### 3.3 Other input data

In addition to the land-use and climate data, most models use additional input data to run their future and past simulations to estimate changes in biodiversity and ecosystem services. For instance, species occurrence data are an integral part of modelling in several of the biodiversity models (i.e. AIM-biodiversity, MOL, cSAR-iDiv, cSAR-IIASA-ETH, BILBI, InSiGHTS) while some models (i.e. cSAR-iDiv, BILBI) rely on estimates of habitat affinity coefficients (e.g. reductions in species richness in a modified habitat relative to the pristine habitat) from the PREDICTS. In DGVM models (i.e. LPJ-GUESS, LPJ, CABLE), atmospheric CO2 concentrations, irrigated fraction and wood harvest estimates are commonly used, while GLOBIO and GLOSP ecosystem services models rely on topography and soil type data for soil erosion measures. A full list of model-specific input data is listed in Table S1.

## 4 Models in BES-SIM

Biodiversity and ecosystem services models at the global scale have increased in number and improved considerably over the last decade, especially with advancement in biodiversity data availability and statistical modelling tools and methods (IPBES, 2016). In order for a model to be included in BES-SIM, it had either to be published in a peer-reviewed journal, or adopt published methodologies, with modifications made to modelling sufficiently documented and accessible for review (Table S2). Sixteen models participated in BES-SIM (Appendix 1, details on modelling methods can be found in Table S2). These models were mainly grouped into four classes: species-based, community-based, and ecosystem-based models of biodiversity, and models of ecosystem functions and services. The methodological approaches, the taxonomic or functional groups, the spatial resolution and the output metrics differ across models (Appendix 1). All sixteen models are spatially explicit with 15 of them using land-use data as an input,12 of them also requiring climate data. We also used one model (BIOMOD2) to assess uncertainty of climate range projections without the use of land-use data.

### 4.1 Species-based models of biodiversity

Species-based models aim to predict historical, current, and future potential distribution and abundance of individual species. These can be developed using correlative methods based on species observation and environmental data (Aguirre-Gutiérrez et al., 2013; Guisan and Thuiller, 2005; Guisan and Zimmermann, 2000), as well as expert-based solutions where data limitations exist (Rondinini et al., 2011). Depending on the methodologies employed and the ecological aspects modelled, they can be known as species distribution models, ecological niche models, bioclimatic envelop models and habitat suitability models (Elith and Leathwick, 2009), and they have been used to forecast environmental impacts on species distribution and status.

In BES-SIM, four species-based models were included: AIM-biodiversity, InSiGHTS, MOL and BIOMOD2 (Appendix 1, Table S2). The first three models project individual species distributions across a large number of species by combining projections of climate impacts on species ranges with projections of land-use impacts on species ranges. AIM (Ohashi et al., in prep.) uses Global Biodiversity Information Facility (GBIF) species occurrence data to train statistical models for current land use and climate to project future species distributions. InSiGHTS (Rondinini et al., 2011; Visconti et al., 2016) and MOL (Jetz et al., 2007; Merow et al., 2013) both rely on expert-based range maps as a baseline. INSIGHTS and MOL used an hierarchical approach with two steps: first, a statistical model trained on current species ranges is used to assess future climate suitability within species ranges; second, an expert-based model detailing associations between species and habitat types is used to assess the impacts of land-use in the climate suitable portion of the species range. BIOMOD2 (Thuiller, 2004; Thuiller et al., 2009) was only used to assess uncertainties in climate-envelope-based projections, and was not included in the model inter comparison with other models incorporating the impacts of land-use change (see section 7. Uncertainties).

### 4.2 Community-based models of biodiversity

Community-based models predict the assemblage of species using environmental data and assess changes in community composition through species presence and abundance (D’Amen et al., 2017). Output variables of community-based models include assemblage-level metrics such as the proportion of species persisting in a landscape, mean species abundances, and compositional similarity relative to a baseline (typically corresponding to a pristine landscape). Three models in BES-SIM (cSAR-iDiv, cSAR-IIASA-ETH, BILBI) rely on versions of the species-area relationship (SAR) to estimate the proportion of species persisting in human-modified habitats relative to native habitat, while three models (PREDICTS, GLOBIO Aquatic, GLOBIO Terrestrial) estimate a range of assemblage-level metrics based on correlative relationships between biodiversity responses and pressure variables (Appendix 1).

Both the cSAR-iDiv (Martins and Pereira, 2017) and the cSAR-IIASA-ETH (Chaudhary et al., 2015) models are based on the countryside species-area relationship (cSAR), which uses habitat affinities to weight the areas of the different habitats in a landscape. The habitat affinities are calibrated from field studies by calculating the change in species richness in a modified habitat relative to the native habitat. The habitat affinities of the cSAR-iDiv model are estimated from the PREDICTS dataset (Hudson et al., 2014) while the habitat affinities of the cSAR-IIASA-ETH come from a previously published database of studies (Chaudhary et al., 2015). The cSAR-iDiv model considers two functional species groups (forest species and non-forest species) for one taxonomic group (birds) while the cSAR-IIASA-ETH uses a single functional group for multiple taxonomic groups (amphibians, birds, mammals, plants and reptiles).

BILBI (Hoskins et al., in prep.; Ferrier et al., 2004, 2007) couples application of the species-area relationship with correlative statistical modelling of continuous patterns of spatial turnover in the species composition of communities as a function of environmental variation. Through space-for-time projection of compositional turnover, this coupled model enables the effects of both climate change and habitat modification to be considered in estimating the proportion of species persisting (in this study for vascular plant species globally).

PREDICTS (Newbold et al., 2016; Purvis et al., 2018) uses a hierarchical mixed-effects framework to model how a range of site-level biodiversity metrics respond to land use and related pressures, using a global database of 767 studies, including over 32,000 sites and 51,000 species. GLOBIO (Alkemade et al., 2009; Janse et al., 2015; Schipper et al., 2016) is an integrative modelling framework for aquatic and terrestrial biodiversity that builds upon correlative relationships between biodiversity intactness and pressure variables, established with meta-analyses of biodiversity monitoring data retrieved from the literature.

### 4.3 Ecosystem-based model of biodiversity

The Madingley model (Harfoot et al., 2014b) is a mechanistic individual-based model of ecosystem structure and function. It encodes a set of fundamental ecological principles to model how individual heterotrophic organisms with a body size greater than 10 μg that feed on other living organisms interact with each other and with their environment. The model is general in the sense that it applies the same set of principles for any ecosystem to which it is applied, and is applicable across scales from local to global. To capture the ecology of all organisms, the model adopts a functional trait based approach with organisms characterised by a set of categorical traits (feeding mode, metabolic pathway, reproductive strategy and movement ability), as well as continuous traits (juvenile, adult and current body mass). Properties of ecological communities emerge from the interactions between organisms, influenced by their environment. The functional diversity of these ecological communities can be calculated as well as the dissimilarity over space or time between communities (Table S2).

### 4.4 Models of ecosystem functions and services

In order to measure ecosystem functions and services, three Dynamic Global Vegetation Models (DGVMs) (i.e., LPJ-GUESS, LPJ, CABLE) and three ecosystem services models (i.e., InVEST, GLOBIO, GLOSP) were engaged in this model intercomparison. The DGVMs are process-based models that simulate responses of potential natural vegetation and associated biogeochemical and hydrological cycles to changes in climate and atmospheric CO2 and disturbance regime (Prentice et al., 2007). Processes in anthropogenically managed land (crop, pasture and managed forests) are also increasingly being accounted for (Arneth et al., 2017). DGVMs can project changes in future ecosystem state and functioning, and habitat structure; however, they are limited in capturing species-level biodiversity change because vegetation is represented by a small number of plant functional types (PFTs) (Bellard et al., 2012; Thuiller et al., 2013).

The InVEST (Sharp et al., 2014) suite includes 18 models that maps and measures the flow and value of ecosystem goods and services across a land or a seascape, based on biophysical processes of the structure and function of ecosystems, accounting for both supply and demand. The GLOBIO model (Alkemade et al., 2009, 2014; Schulp et al., 2012) estimates ecosystem services based on outputs from the IMAGE model (Stehfest et al., 2014), the global hydrological model PCRaster Global Water Balance (PCR-GLOBWB, van Beek et al., 2011), and the Global Nutrient Model (Beusen et al., 2015). It is based on correlative relationships between ecosystem functions and services and particular environmental variables (mainly land use), quantified based on literature data. Finally, the GLOSP (Guerra et al., 2016) is a 2D model that estimates the level of global and local soil erosion and protection using the Universal Soil Loss Equation.

## 5 Output metrics

Given the diversity of modelling approaches, a wide range of biodiversity and ecosystem services metrics can be produced by the model set (Table S2). For the biodiversity model intercomparison analysis, three main categories of common output metrics were reported over time: extinctions as absolute change in species richness (N) or as proportional species richness change (P); abundance-based intactness (I); and mean proportional change in suitable habitat extent across species (H) (Table 4). These metrics were calculated at two scales: local or grid cell (α) and regional or global (γ). Some models only provided α values while others provided both α and γ values (Table 4). For the models that can project γ metrics, both regional-γ for each IPBES regions and a global-γ were reported. Absolute changes in species richness and proportional species richness change are interrelated and may be calculated from reporting species richness over time, as N_t_=S_t_-S_t0_ and P=N_t_/S_t0_, where S_t_ is the number of species at time *t*. Most models reported one or both types of species richness metrics (Table 4). Intactness, which can be estimated in several ways, refers to the difference between the current community composition and the inferred original state in the native vegetation. This metric is available only for two community-based models (i.e., GLOBIO and PREDICTS). The habitat change (H) was calculated from averaging across species occurring in the unit of analysis (grid cell, region, or globe) the changes in the suitable habitat extent of each species relative to a baseline, i.e. (E_i,t_-E_i,t0_)/E_i,t0_, where E_i,t_ is the suitable habitat extent of species *i* at time *t* within the unit of analysis. It can be reported for species-level models (i.e. AIM-biodiversity, InSiGHTS, MOL) (Table 4). The baseline year, t_0_, used to calculate changes for the extinction and habitat extent metrics was the first year of the simulation (in most cases t_0_=1900, see Table 5).

**Table 4:** Selected output indicators for inter-comparison of biodiversity and ecosystems models. For species diversity change, both proportional changes in species richness (P) and absolute changes (N) are reported. Some models project the α metrics at the level of the grid cell (e.g. species-based and SAR based community models) while others average the local values of the metrics across the grid cell weighted by the area of the different habitats in the cell (e.g. PREDICTS, GLOBIO).

| BES-SIM model | Local scale species diversity change (Pα and Nα) | Subregional and global scale species diversity change (Pγ and Nγ) | Abundance-based intactness (Iα) | Local and global mean habitat extent change (Hα and Hγ) |
| --- | --- | --- | --- | --- |
| **Species-based models of biodiversity** |  |  |  |  |
| AIM-biodiversity | * | * |  | * |
| InSiGHTS | * | * |  | * |
| MOL | * | * |  | * |
| **Community-based models of biodiversity** |  |  |  |  |
| cSAR-iDiv | * | * |  |  |
| cSAR-IIASA-ETH | * | * |  |  |
| BILBI |  | * |  |  |
| PREDICTS | * |  | * |  |
| GLOBIO - Aquatic |  |  | * |  |
| GLOBIO - Terrestrial |  |  | * |  |
| **Ecosystems-based model of biodiversity** |  |  |  |  |
| Madingley |  |  | * |  |

**Table 5:** Scenario (forcing data) for models in BES-SIM.

|  |  | Future Land-Use Change or Climate<br>(2050) |  |  |
| --- | --- | --- | --- | --- |
| BES-SIM model | Historical | Land use only, climate held<br>constant at 2015<br>(SSP1, SSP3, SSP5) | Climate change only, land use<br>held constant at 2015<br>(RCP2.6, RCP6.0, RCP8.5) | Land use and climate<br>(SSP1xRCP2.6,<br>SSP3xRCP6.0, SSP5xRCP8.5) |
| **Species-based models of biodiversity** |  |  |  |  |
| AIM-biodiversity | * | * | * | * |
| InSiGHTS | * | * | * | * |
| MOL |  | * | * | * |
| **Community-based models of biodiversity** |  |  |  |  |
| cSAR-iDiv | * | * |  |  |
| cSAR-IIASA-ETH | * | * |  |  |
| BILBI | * | * |  | * |
| PREDICTS | * | * |  |  |
| GLOBIO - Aquatic |  |  |  | * |
| GLOBIO - Terrestrial |  | * | * | * |
| **Ecosystems-based model of biodiversity** |  |  |  |  |
| Madingley | * |  |  | * |
| **Models of ecosystem functions and services** |  |  |  |  |
| LPJ-GUESS | * | * | * | * |
| LPJ | * | * | * | * |
| CABLE | * | * | * | * |
| GLOBIO-ES |  |  |  | * |
| InVEST |  |  |  | * |
| GLOSP |  |  |  | * |

For ecosystem functions and services, each model’s output metrics were mapped onto the new classification of Nature’s Contributions to People (NCP) published by the IPBES scientific community (Díaz et al., 2018). Among the 18 possible NCPs, the combination of models participating in BES-SIM were able to provide measures for 10 NCPs, including regulating metrics on pollination (e.g., proportion of agricultural lands whose pollination needs are met), climate (e.g., vegetation carbon, total carbon uptake and loss), water quantity (e.g., monthly runoff), water quality (e.g., nitrogen and phosphorus leaching, algal blooms), soil protection (e.g., erosion risk), hazards (e.g., costal resilience, flood risk) and detrimental organisms (e.g. fraction of cropland potentially protected by the natural pest, relative to all available cropland), and material metrics on bioenergy (e.g. bioenergy-crop production), food and feed (e.g. total crop production) and materials (e.g. wood harvest) (Table 6). Some of these metrics require careful interpretation in the context of NCPs (e.g., an increase in flood risk can be caused by climate change and/or by a reduction of the capacity of ecosystems to reduce flood risk) and additional translation of increasing or declining measures of ecosystem functions and services (e.g., food and feed, water quantity) into contextually relevant information (i.e., positive or negative impacts) on human well-being and quality of life. Given disparity of metrics across models within each NCP category, names and units of the metrics are listed in Table 6 with definitions and methods provided in Table S3.

**Table 6:** Selected output indicators for inter-comparison of ecosystem functions and services models, categorized based on the classification of Nature’s Contributions to People (Díaz et al., 2018).

| BES-SIM model | NCP 2.<br>Pollination and dispersal of seeds and other propagules | NCP 4.<br>Regulation of climate | NCP 6.<br>Regulation of freshwater quantity, location and timing | NCP 7.<br>Regulation of freshwater and coastal water quality | NCP 8.<br>Formation, protection and decontamination of soils and sediments | NCP 9.<br>Regulation of hazards and extreme events | NCP 10.<br>Regulation of detrimental organisms and biological processes | NCP 11.<br>Energy | NCP 12.<br>Food and feed | NCP 13.<br>Materials, companionship and labor |
| --- | --- | --- | --- | --- | --- | --- | --- | --- | --- | --- |
| LPJ-GUESS |  | Total carbon<br>Vegetation carbon | Monthly runoff | Nitrogen leaching |  |  |  | Bioenergy-crop production | Harvested carbon in croplands that are used for food production | Wood harvest (LUH2 extraction) |
| LPJ |  | Total carbon<br>Vegetation carbon | Monthly runoff |  |  |  |  |  |  |  |
| CABLE |  | Total carbon<br>Vegetation carbon | Monthly runoff,<br>Total runoff |  |  |  |  |  | Above ground carbon removed from cropland and pastures as a result of harvest and grazing | Wood harvest |
| GLOBIO-ES | Fraction of cropland potentially pollinated, relative to all available cropland | Total carbon | Water scarcity index | Nitrogen in water<br>Phosphorus in water | Erosion protection: fraction with low risk relative to the area that needs protection | Flood risk: number of people exposed to river flood risk | Pest control: Fraction of cropland potentially protected, relative to all available cropland |  | Total crop production<br>Total grass production |  |
| InVEST | Proportion of agricultural lands whose pollination needs are met |  |  | Nitrogen export<br>Nitrogen export*capita |  | Coastal vulnerability<br>Coastal vulnerability*capita |  |  | Caloric production per hectare on the current landscape for each crop type |  |
| GLOSP |  |  |  |  | Soil protection |  |  |  |  |  |

## 6 Core simulations

The simulations for BES-SIM required a minimum of two outputs from the modelling teams: present (2015) and future (2050). Additional, a past projection (1900) and a farther future projection (2070) were also provided by several modelling teams. Some models projected further into the past and also at multiple time points from the past to the future (Table 5). Models that simulated a continuous time-series of climate change (and land-use change) impacts provided 20-year averages around these mid-points to account for inter-annual variability. The models ran simulations at their original spatial resolutions (Appendix 1), and upscaled results to one-degree grid cells using arithmetic means. In order to provide global or regional averages of the α or grid cell metrics, the arithmetic mean values across the cells of the globe or a region were calculated, as well as percentiles of those metrics. Both, one-degree rasters and a table with values for each IPBES region and the globe were provided by each modelling team for each output metric.

To measure the individual and synergistic impacts of land use and climate change on biodiversity and ecosystem services, models accounting for both types of drivers were run three times: with land-use change, with climate change only, and with both drivers combined. For instance, to measure the impact of land use alone, the projections into 2050 were obtained while retaining climate data constant from present (2015) to the future (2050). Similarly, to measure the impact of climate change alone, the climate projections into 2050 (or 2070) were obtained while retaining the land-use data constant from present (2015) to the future (2050). Finally, to measure the impact of land use and climate change combined, models were run using projections of both land use and climate change into 2050 (or 2070). When backcasting to 1900, for the models that used ISI-MIP 2a IPSL climate dataset, random years from 1951 to 1960 were selected to fill the gap in climate input for years 1901 to 1950. The models (i.e., InSiGHTS, BILBI) that used WorldClim dataset did not simulate climate scenarios for the past projections given the gap in climate input before 1960.

## 7 Uncertainties

Reporting uncertainty is a critical component of model intercomparison exercises (IPBES, 2016). Within BES-SIM, uncertainties were explored in two ways: (1) each model had to report the mean values of its metrics, and where possible the 25^th^, 50^th^, and 75^th^ percentiles based on different model parameterizations; and when combining the data provided by the different models, the average and the standard deviation of the common metrics were calculated (e.g., intermodel average and standard deviation of Pγ); (2) the BIOMOD model was used in assessing the uncertainty in modelling changes in species ranges arising from using different RCP scenarios, different GCMs, a suite of species distribution modelling algorithms (e.g., random forest, logistic regression) and different species dispersal hypotheses.

In the intercomparison analysis, we will conduct a comprehensive uncertainty analysis based on a variance partitioning approach on the outputs provided by the models of biodiversity. This will allow us to highlight uncertainties arising from the land use (SSPs), the climate (RCPs and GCMs), and, where relevant, the different taxa.

## 8 Discussion

This manuscript lays out the context, motives, processes, and approaches taken for a scenario-based intercomparison of biodiversity and ecosystem service models (BES-SIM). This model intercomparison initiative aims to provide scientifically rigorous information to the IPBES and its ongoing and future assessments, the CBD and its strategic plans and conservation goals, and other relevant stakeholders on the expected status and trends of biodiversity and ecosystem services using a suite of metrics from a range of global models. The resulting outputs will include the analyses on the past, present and future impacts of land-use change, climate change and other drivers as embodied in a range of human development scenarios, coupled with associated climate projections. The model intercomparison analyses will put the future in the context of the past and the present.

The existing SSP and RCP scenarios provided a consistent set of past and future projections of two major drivers of terrestrial and freshwater biodiversity change – land use and climate. However, we acknowledge that these projections have certain limitations. These include limited inclusion of biodiversity-specific policies in the storylines (only the SSP1 baseline emphasises additional biodiversity policies) (O’Neill et al., 2016; Rosa et al., 2017), coarse spatial resolution, and land-use classes that are not sufficiently detailed to fully capture the response of biodiversity to land-use change (Harfoot et al., 2014a; Titeux et al., 2016, 2017). The heterogeneity of models and their methodological approaches, as well as additional harmonization of metrics of ecosystem functions and services (Tables 6, S3) are areas for further work. In the future, it will be also important to capture the uncertainties associated with input data, with a focus on uncertainty in land-use and climate projections resulting from differences among IAMs and GCMs on each scenario (Popp et al., 2017). The gaps identified through BES-SIM and future directions for research and modelling will be published with analyses of the results on the model intercomparison and on individual models.

As a long-term perspective, BES-SIM is expected to provide critical foundation and insights for the ongoing development of nature-centred, multiscale Nature Futures scenarios (Rosa et al., 2017). Catalysed by the IPBES Expert Group on Scenarios and Models, this new scenarios and modelling framework will shift traditional ways of forecasting impacts of society on nature to more integrative, biodiversity-centred visions and pathways. A future round of BES-SIM could use these biodiversity-centred storylines to project dynamics of biodiversity and ecosystem services and associated consequences for human well-being and socio-economic development. This will help policymakers and practitioners to collectively identify pathways for sustainable futures based on alternative biodiversity management approaches and assist researchers in incorporating the role of biodiversity on socio-economic scenarios.

## 9. Code and data availability

The output data from this model intercomparison will be downloadable from the website of the IPBES Expert Group on Scenarios and Models in the future (https://www.ipbes.net/deliverables/3c-scenarios-and-modelling). The LUH2 land-use data used for model runs are available on http://luh.umd.edu/data.shtml. The climate datasets used in BES-SIM can be downloaded from the respective websites (https://www.isimip.org/outputdata/, http://worldclim.org/version1)

## Footnotes

Author contributions. All authors co-designed the study under the coordination of Henrique M. Pereira, Rob Alkemade, Paul Leadley and Isabel M.D. Rosa. HyeJin Kim prepared the manuscript with contributions from all co-authors.

## Acknowledgements

HJK, ISM, FW, CG and HMP are supported by the German Centre for integrative Biodiversity Research (iDiv) Halle-Jena-Leipzig, funded by the German Research Foundation (FZT 118). IMDR received funding from the European Union’s Horizon 2020 research and innovation programme under the Marie Sklodowska-Curie grant agreement No 703862. PL was supported by the LabEx BASC supported by the French “Investment ďAvenif” program (grant ANR-11-LABX-0034). GCH and LPC gratefully acknowledge the support of DOE-SciDAC program. AA, AK, BQ and PA acknowledge support from the Helmholtz Association and its ATMO Programme. Support was also provided by the EU FP7 project LUC4C. AP, ADP and SLLH are supported by the Natural Environment Research Council U.K. (grant number NE/M014533/1) and by a DIF grant from the Natural History Museum. RCK and RS were supported by private gifts to the Natural Capital Project. DL, FDF, PH, and MO are supported by the project IS-WEL-Integrated Solutions for Water, Energy and Land funding from Global Environmental Facility, Washington, USA, coordinated by United Nations Industrial Development Organization (UNIDO), UNIDO Project No. 140312. FDF and MO are supported by the ERC SYNERGY grant project IMBALANCE-P-Managing Phosphorous limitation in a nitrogen-saturated Anthropocene, funding from European Commission, European Research Council Executive Agency, grant agreement No. 610028. DL and PH were supported by the project SIGMA-Stimulating Innovation for Global Monitoring of Agriculture and its Impact on the Environment in support of GEOGLAM, funding from the European Union’s FP7 research and innovation programme under the Environment area, grant agreement No. 603719. TH, HO, AH, SF, TM and KT are supported by the Global Environmental Research (S-14) of the Ministry of the Environment of Japan. TH, SF and KT are supported by Environment Research and Technology Development Fund 2-1702 of the Environmental Restoration and Conservation Agency of Japan and JSPS Overseas Research Fellowships. MH was supported by a KR Rasmussen Foundation grant “Modelling the Biodiversity Planetary Boundary and Embedding Results into Policy”. VH acknowledges support from the Earth Systems and Climate Change Hub, funded by the Australian Government’s National Environmental Science Program. CM acknowledges funding from NSF Grant DEB1565046. Finally, we also thank the following organizations for funding the workshops: PBL Netherland Environment Assessment Agency, UNESCO (March 2016), iDiv German Centre for integrative Biodiversity Research (October 2016, October 2017) and Zoological Society of London (January 2018).

## Appendix 1. Description of biodiversity and ecosystem functions and services models in BES-SIM.

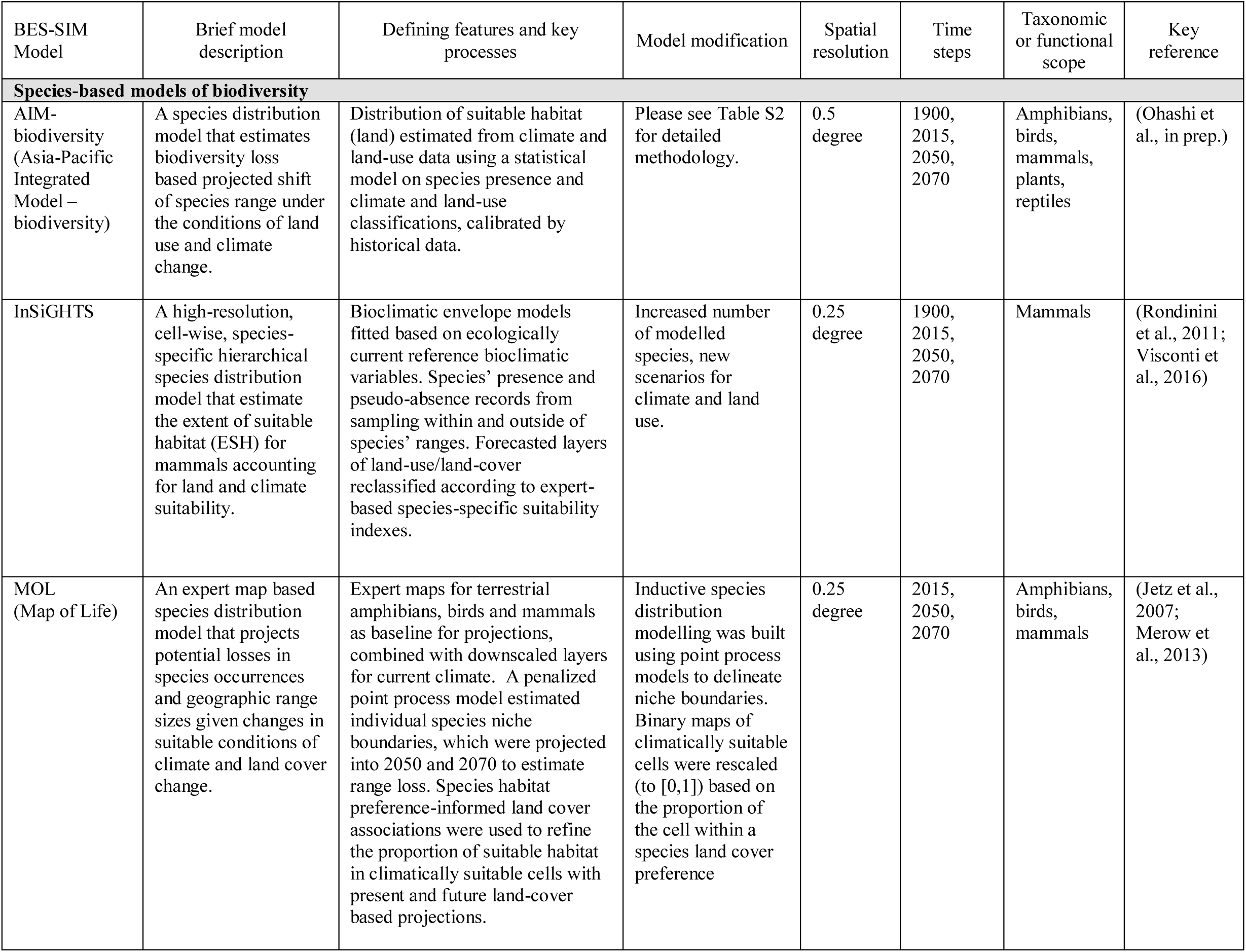
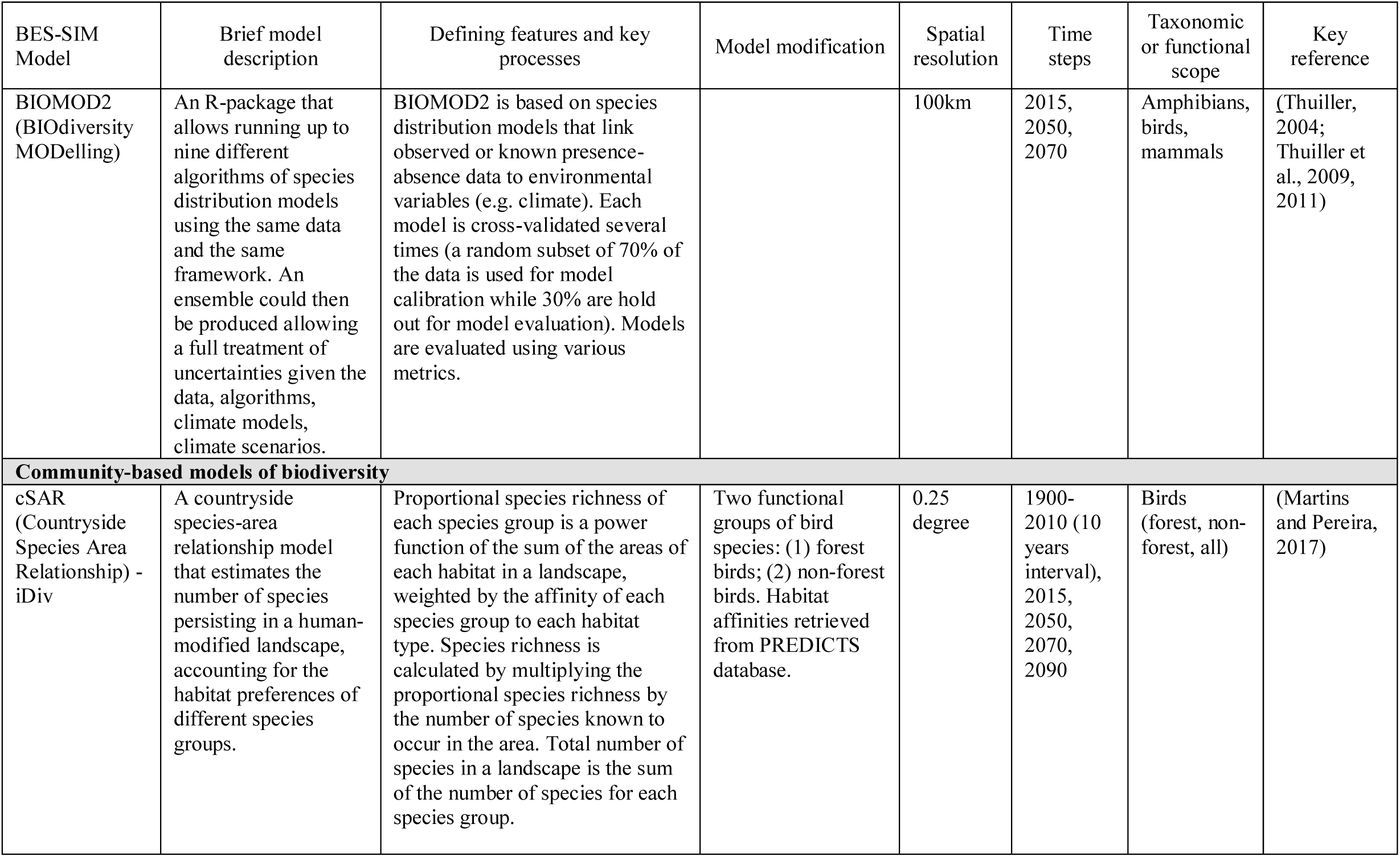
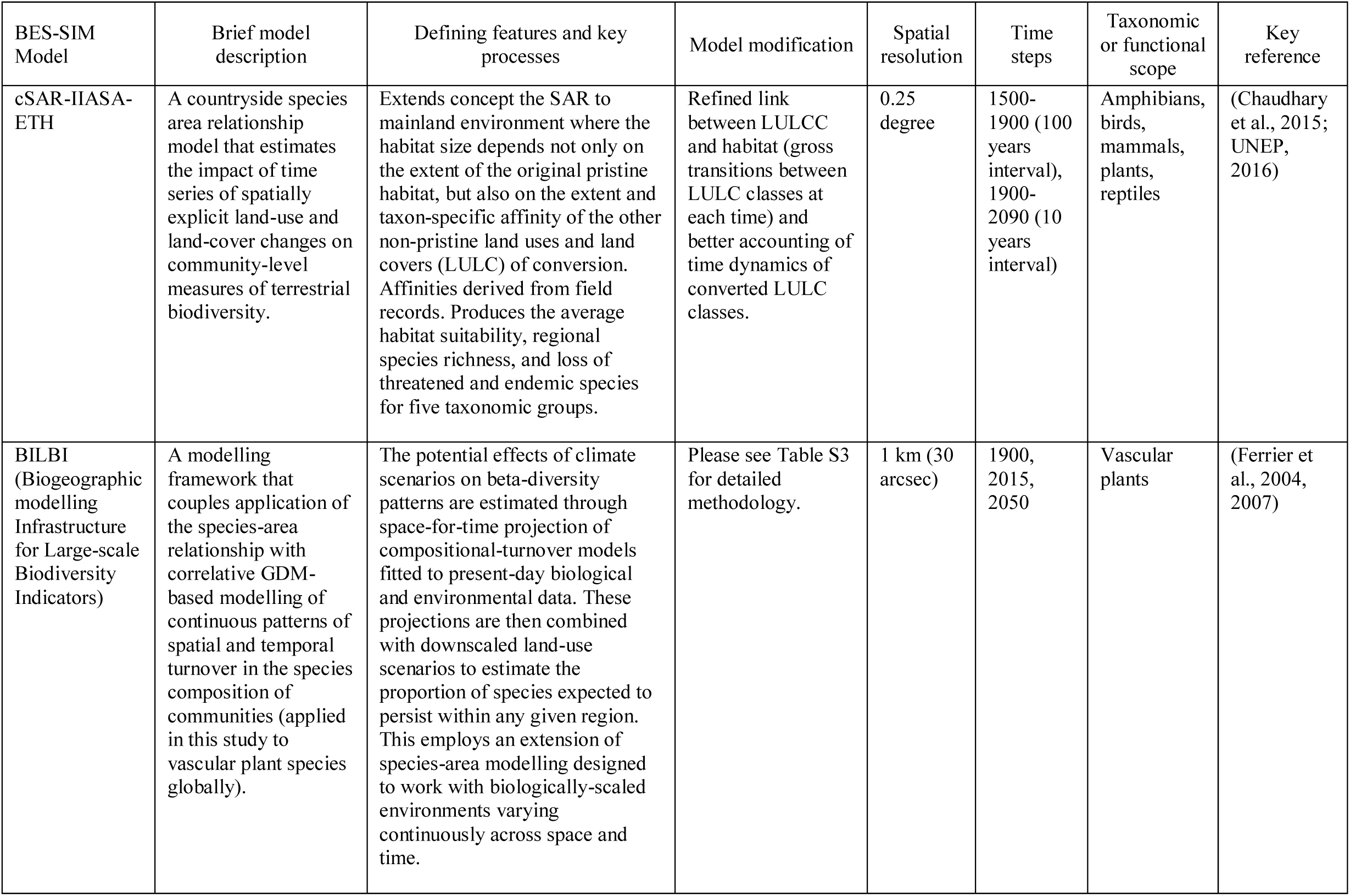
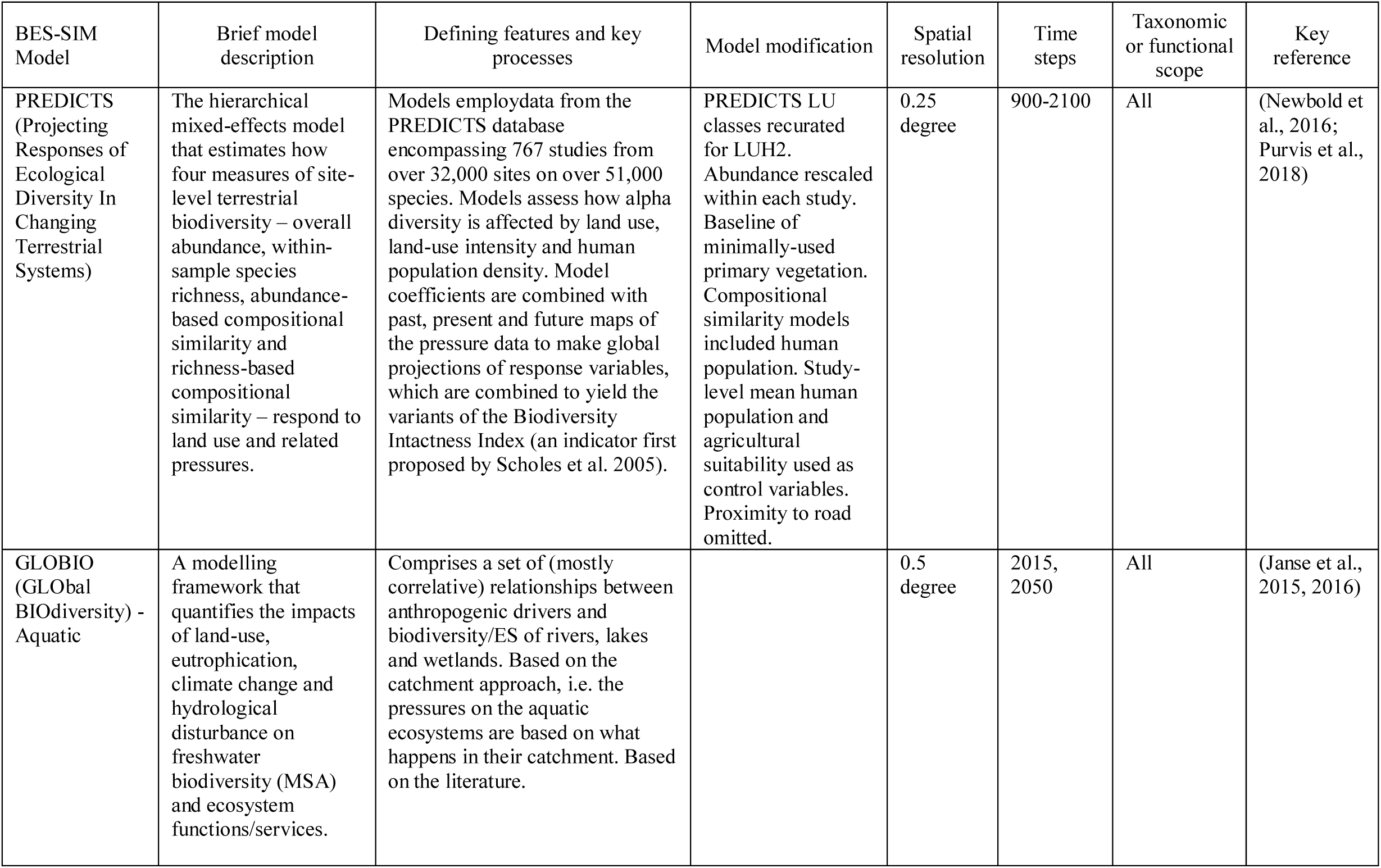
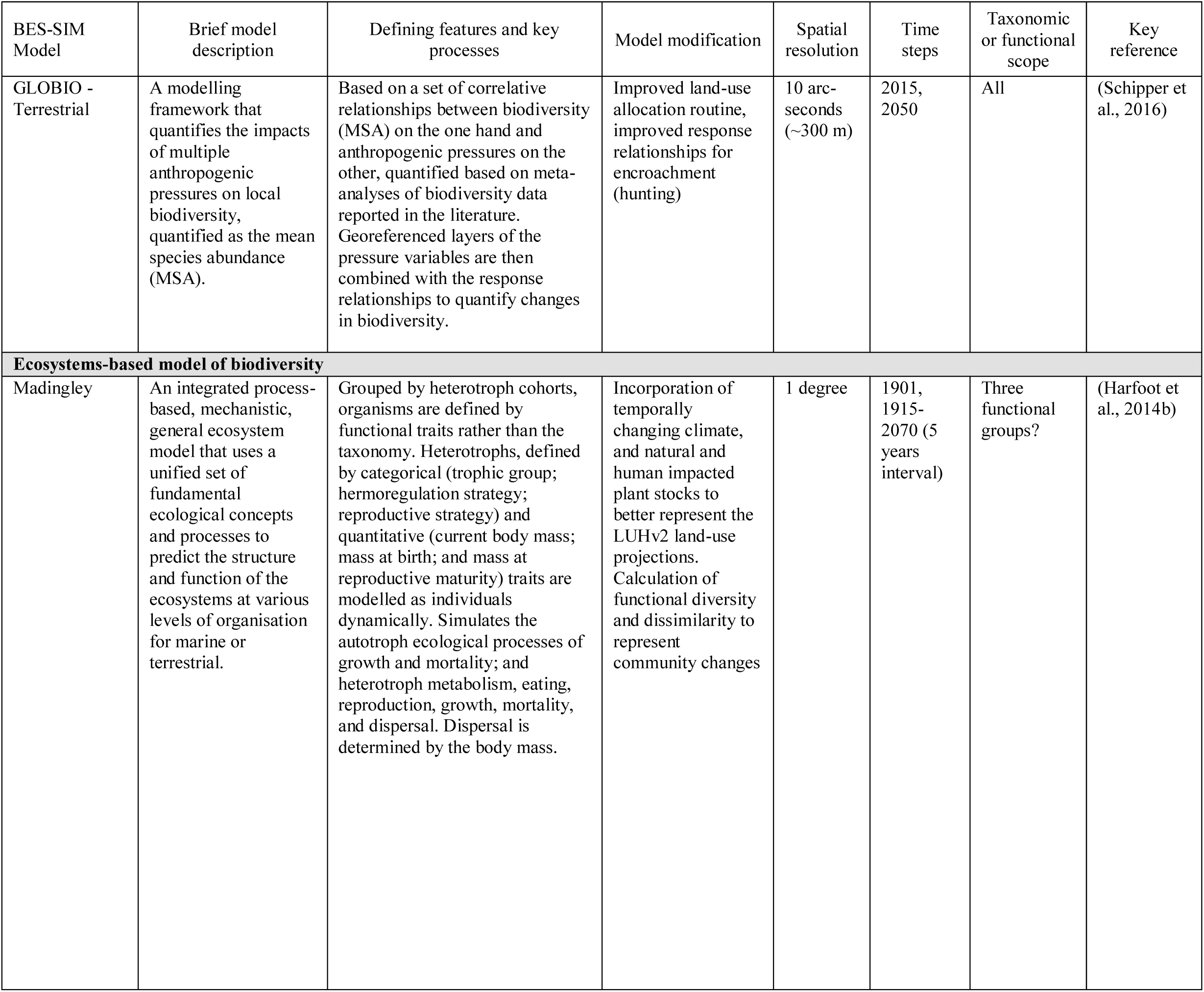
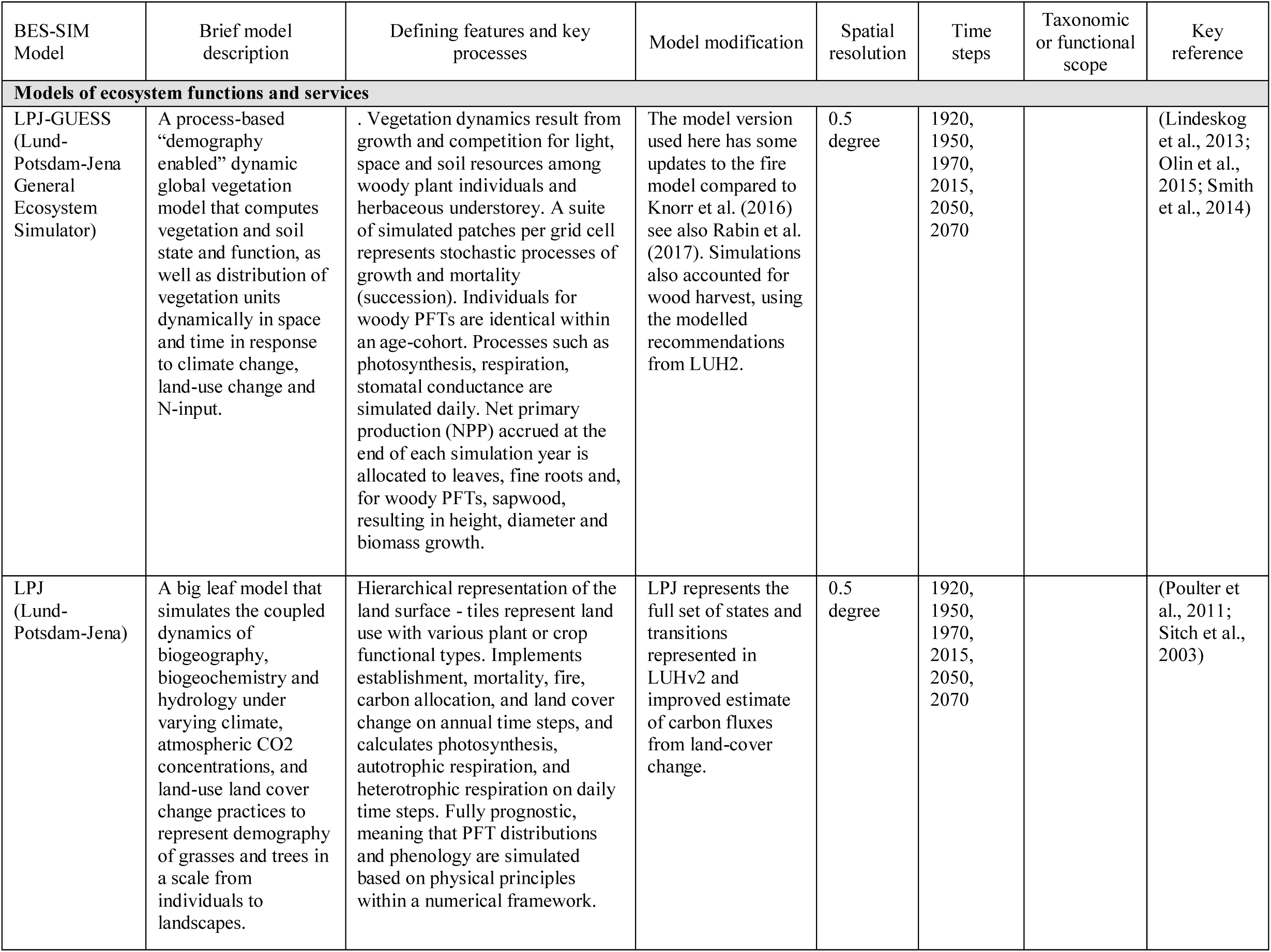
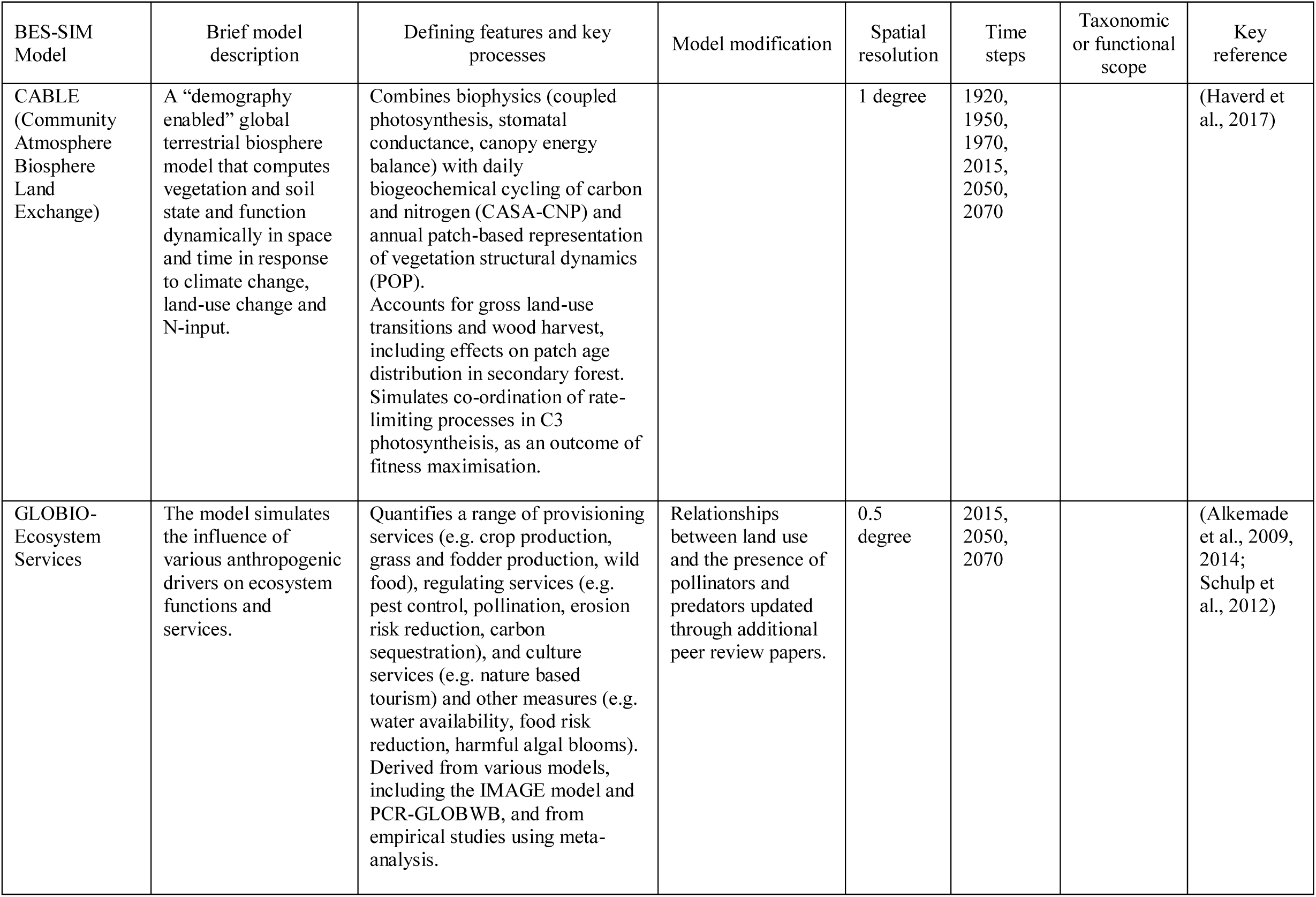
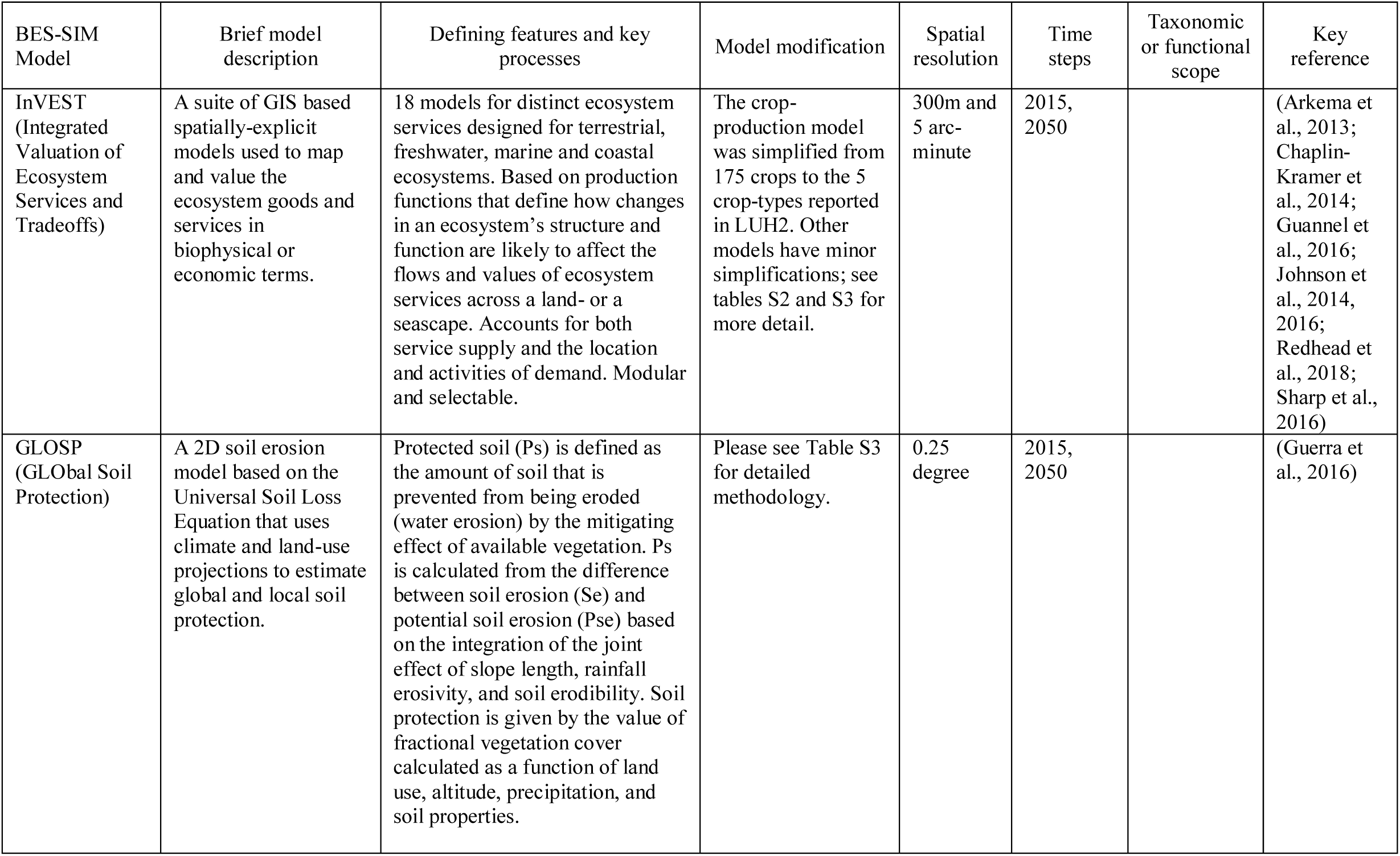

## Appendix 2. Acronyms

AIM: Asia-pacific Integrated Model
BES-SIM: Biodiversity and Ecosystem Services Scenario-based Intercomparison of Models
BIOMOD: BIOdiversity MODelling
BILBI: Biogeographic modelling Infrastructure for Large-scale Biodiversity Indicators
CABLE: Community Atmosphere Biosphere Land Exchange
CMIP: Climate Model Inter-comparison Project
cSAR: Countryside Species Area Relationship
DGVM: Dynamic Global Vegetation Model
ESM: Earth System Models
GBIF: Global Biodiversity Information Facility
GBO: Global Biodiversity Outlooks
GCM: Global Circulation Models
GEO: Global Environmental Outlook
GLOBIO: GLObal BIOdiversity
GLOSP: GLObal Soil Protection
IAM: Integrated Assessment Models
IMAGE: Integrated Model to Assess the Global Environment
InVEST: Integrated Valuation of Ecosystem Services and Tradeoffs
IPBES: Intergovernmental Science-Policy Platform on Biodiversity and Ecosystem Services
IPCC: Intergovernmental Panel on Climate Change
IPSL-CM5A-LR: Institut Pierre-Simon Laplace-Climate Model 5A-Low Resolution
ISI-MIP: Inter-Sectoral Impact Model Intercomparison Project
LPJ: Lund-Potsdam-Jena
LPJ-GUESS: Lund-Potsdam-Jena General Ecosystem Simulator
LUH2: Land Use Harmonization Project version 2
MA: Millennium Ecosystem Assessment
MAgPIE: The Model of Agricultural Production and its Impact on the Environment
MIP: Model Intercomparison Project
MOL: Map of Life
NCP: Nature’s Contributions to People
REMIND: Regionalized Model of Investments and Development
PREDICTS: Projecting Responses of Ecological Diversity In Changing Terrestrial Systems
RCM: Regional Climate Models
RCPs: Representative Concentration Pathways
SAR: Species Area Relationship
SR: Species Richness
SSPs: Shared Socio-economic Pathways

## References

Aguirre-Gutiérrez, J., Carvalheiro, L. G., Polce, C., van Loon, E. E., Raes, N., Reemer, M. and Biesmeijer, J. C.: Fit-for-Purpose: Species Distribution Model Performance Depends on Evaluation Criteria – Dutch Hoverflies as a Case Study, edited by M. G. Chapman, PLoS ONE, 8(5), e63708, doi:10.1371/journal.pone.0063708, 2013.

Alkemade, R., van Oorschot, M., Miles, L., Nellemann, C., Bakkenes, M. and ten Brink, B.: GLOBIO3: A Framework to Investigate Options for Reducing Global Terrestrial Biodiversity Loss, Ecosystems, 12(3), 374–390, doi:10.1007/s10021-009-9229-5, 2009.

Alkemade, R., Burkhard, B., Crossman, N. D., Nedkov, S. and Petz, K.: Quantifying ecosystem services and indicators for science, policy and practice, Ecol. Indic., 37, 161–162, doi:10.1016/j.ecolind.2013.11.014, 2014.

Arkema, K. K., Guannel, G., Verutes, G., Wood, S. A., Guerry, A., Ruckelshaus, M., Kareiva, P., Lacayo, M. and Silver, J. M.: Coastal habitats shield people and property from sea-level rise and storms, Nat. Clim. Change, 3(10), 913–918, doi:10.1038/nclimate1944, 2013.

Arneth, A., Sitch, S., Pongratz, J., Stocker, B. D., Ciais, P., Poulter, B., Bayer, A. D., Bondeau, A., Calle, L., Chini, L. P., Gasser, T., Fader, M., Friedlingstein, P., Kato, E., Li, W., Lindeskog, M., Nabel, J. E. M. S., Pugh, T. A. M., Robertson, E., Viovy, N., Yue, C. and Zaehle, S.: Historical carbon dioxide emissions caused by land-use changes are possibly larger than assumed, Nat. Geosci., 10(2), 79–84, doi:10.1038/ngeo2882, 2017.

van Beek, L. P. H., Wada, Y. and Bierkens, M. F. P.: Global monthly water stress: 1. Water balance and water availability: GLOBAL MONTHLY WATER STRESS, 1, Water Resour. Res., 47(7), doi:10.1029/2010WR009791, 2011.

Bellard, C., Bertelsmeier, C., Leadley, P., Thuiller, W. and Courchamp, F.: Impacts of climate change on the future of biodiversity: Biodiversity and climate change, Ecol. Lett., 15(4), 365–377, doi:10.1111/j.1461-0248.2011.01736.x, 2012.

Beusen, A. H. W., Van Beek, L. P. H., Bouwman, A. F., Mogollón, J. M. and Middelburg, J. J.: Coupling global models for hydrology and nutrient loading to simulate nitrogen and phosphorus retention in surface water – description of IMAGE–GNM and analysis of performance, Geosci. Model Dev., 8(12), 4045–4067, doi:10.5194/gmd-8-4045-2015, 2015.

Cardinale, B. J., Duffy, J. E., Gonzalez, A., Hooper, D. U., Perrings, C., Venail, P., Narwani, A., Mace, G. M., Tilman, D., Wardle, D. A., Kinzig, A. P., Daily, G. C., Loreau, M., Grace, J. B., Larigauderie, A., Srivastava, D. S. and Naeem, S.: Biodiversity loss and its impact on humanity, Nature, 486(7401), 59–67, doi:10.1038/nature11148, 2012.

Chaplin-Kramer, R., Dombeck, E., Gerber, J., Knuth, K. A., Mueller, N. D., Mueller, M., Ziv, G. and Klein, A.-M.: Global malnutrition overlaps with pollinator-dependent micronutrient production, Proc. R. Soc. B Biol. Sci., 281(1794), 20141799–20141799, doi:10.1098/rspb.2014.1799, 2014.

Chaudhary, A., Verones, F., de Baan, L. and Hellweg, S.: Quantifying Land Use Impacts on Biodiversity: Combining Species–Area Models and Vulnerability Indicators, Environ. Sci. Technol., 49(16), 9987–9995, doi:10.1021/acs.est.5b02507, 2015.

D’Amen, M., Rahbek, C., Zimmermann, N. E. and Guisan, A.: Spatial predictions at the community level: from current approaches to future frameworks: Methods for community-level spatial predictions, Biol. Rev., 92(1), 169–187, doi:10.1111/brv.12222, 2017.

Díaz, S., Pascual, U., Stenseke, M., Martín-López, B., Watson, R. T., Molnár, Z., Hill, R., Chan, K. M. A., Baste, I. A., Brauman, K. A., Polasky, S., Church, A., Lonsdale, M., Larigauderie, A., Leadley, P. W., van Oudenhoven, A. P. E., van der Plaat, F., Schröter, M., Lavorel, S., Aumeeruddy-Thomas, Y., Bukvareva, E., Davies, K., Demissew, S., Erpul, G., Failler, P., Guerra, C. A., Hewitt, C. L., Keune, H., Lindley, S. and Shirayama, Y.: Assessing nature’s contributions to people, Science, 359(6373), 270–272, doi:10.1126/science.aap8826, 2018.

Dufresne, J.-L., Foujols, M.-A., Denvil, S., Caubel, A., Marti, O., Aumont, O., Balkanski, Y., Bekki, S., Bellenger, H., Benshila, R., Bony, S., Bopp, L., Braconnot, P., Brockmann, P., Cadule, P., Cheruy, F., Codron, F., Cozic, A., Cugnet, D., de Noblet, N., Duvel, J.-P., Ethé, C., Fairhead, L., Fichefet, T., Flavoni, S., Friedlingstein, P., Grandpeix, J.-Y., Guez, L., Guilyardi, E., Hauglustaine, D., Hourdin, F., Idelkadi, A., Ghattas, J., Joussaume, S., Kageyama, M., Krinner, G., Labetoulle, S., Lahellec, A., Lefebvre, M.-P., Lefevre, F., Levy, C., Li, Z. X., Lloyd, J., Lott, F., Madec, G., Mancip, M., Marchand, M., Masson, S., Meurdesoif, Y., Mignot, J., Musat, I., Parouty, S., Polcher, J., Rio, C., Schulz, M., Swingedouw, D., Szopa, S., Talandier, C., Terray, P., Viovy, N. and Vuichard, N.: Climate change projections using the IPSL-CM5 Earth System Model: from CMIP3 to CMIP5, Clim. Dyn., 40(9–10), 2123–2165, doi:10.1007/s00382-012-1636-1, 2013.

Elith, J. and Leathwick, J. R.: Species Distribution Models: Ecological Explanation and Prediction Across Space and Time, Annu. Rev. Ecol. Evol. Syst., 40(1), 677–697, doi:10.1146/annurev.ecolsys.110308.120159, 2009.

Ferrier, S., Powell, G. V. N., Richardson, K. S., Manion, G., Overton, J. M., Allnutt, T. F., Cameron, S. E., Mantle, K., Burgess, N. D., Faith, D. P., Lamoreux, J. F., Kier, G., Hijmans, R. J., Funk, V. A., Cassis, G. A., Fisher, B. L., Flemons, P., Lees, D., Lovett, J. C. and Van Rompaey, R. S. A. R.: Mapping More of Terrestrial Biodiversity for Global Conservation Assessment, BioScience, 54(12), 1101, doi:10.1641/0006-3568(2004)054[1101:MMOTBF]2.0.CO;2, 2004.

Ferrier, S., Manion, G., Elith, J. and Richardson, K.: Using generalized dissimilarity modelling to analyse and predict patterns of beta diversity in regional biodiversity assessment, Divers. Distrib., 13(3), 252–264, doi:10.1111/j.1472-4642.2007.00341.x, 2007.

Fick, S. E. and Hijmans, R. J.: WorldClim 2: new 1-km spatial resolution climate surfaces for global land areas: NEW CLIMATE SURFACES FOR GLOBAL LAND AREAS, Int. J. Climatol., 37(12), 4302–4315, doi:10.1002/joc.5086, 2017.

Frieler, K., Levermann, A., Elliott, J., Heinke, J., Arneth, A., Bierkens, M. F. P., Ciais, P., Clark, D. B., Deryng, D., Döll, P., Falloon, P., Fekete, B., Folberth, C., Friend, A. D., Gellhorn, C., Gosling, S. N., Haddeland, I., Khabarov, N., Lomas, M., Masaki, Y., Nishina, K., Neumann, K., Oki, T., Pavlick, R., Ruane, A. C., Schmid, E., Schmitz, C., Stacke, T., Stehfest, E., Tang, Q., Wisser, D., Huber, V., Piontek, F., Warszawski, L., Schewe, J., Lotze-Campen, H. and Schellnhuber, H. J.: A framework for the cross-sectoral integration of multi-model impact projections: land use decisions under climate impacts uncertainties, Earth Syst. Dyn., 6(2), 447–460, doi:10.5194/esd-6-447-2015, 2015.

Frieler, K., Lange, S., Piontek, F., Reyer, C. P. O., Schewe, J., Warszawski, L., Zhao, F., Chini, L., Denvil, S., Emanuel, K., Geiger, T., Halladay, K., Hurtt, G., Mengel, M., Murakami, D., Ostberg, S., Popp, A., Riva, R., Stevanovic, M., Suzuki, T., Volkholz, J., Burke, E., Ciais, P., Ebi, K., Eddy, T. D., Elliott, J., Galbraith, E., Gosling, S. N., Hattermann, F., Hickler, T., Hinkel, J., Hof, C., Huber, V., Jägermeyr, J., Krysanova, V., Marcé, R., Müller Schmied, H., Mouratiadou, I., Pierson, D., Tittensor, D. P., Vautard, R., van Vliet, M., Biber, M. F., Betts, R. A., Bodirsky, B. L., Deryng, D., Frolking, S., Jones, C. D., Lotze, H. K., Lotze-Campen, H., Sahajpal, R., Thonicke, K., Tian, H. and Yamagata, Y.: Assessing the impacts of 1.5 °C global warming – simulation protocol of the Inter-Sectoral Impact Model Intercomparison Project (ISIMIP2b), Geosci. Model Dev., 10(12), 4321–4345, doi:10.5194/gmd-10-4321-2017, 2017.

Frischknecht, R., Fantke, P., Tschümperlin, L., Niero, M., Antón, A., Bare, J., Boulay, A.-M., Cherubini, F., Hauschild, M. Z., Henderson, A., Levasseur, A., McKone, T. E., Michelsen, O., i Canals, L. M., Pfister, S., Ridoutt, B., Rosenbaum, R. K., Verones, F., Vigon, B. and Jolliet, O.: Global guidance on environmental life cycle impact assessment indicators: progress and case study, Int. J. Life Cycle Assess., 21(3), 429–442, doi:10.1007/s11367-015-1025-1, 2016.

Fujimori, S., Hasegawa, T., Masui, T., Takahashi, K., Herran, D. S., Dai, H., Hijioka, Y. and Kainuma, M.: SSP3: AIM implementation of Shared Socioeconomic Pathways, Glob. Environ. Change, 42, 268–283, doi:10.1016/j.gloenvcha.2016.06.009, 2017.

Guannel, G., Arkema, K., Ruggiero, P. and Verutes, G.: The Power of Three: Coral Reefs, Seagrasses and Mangroves Protect Coastal Regions and Increase Their Resilience, edited by C. N. Bianchi, PLOS ONE, 11(7), e0158094, doi:10.1371/journal.pone.0158094, 2016.

Guerra, C. A., Maes, J., Geijzendorffer, I. and Metzger, M. J.: An assessment of soil erosion prevention by vegetation in Mediterranean Europe: Current trends of ecosystem service provision, Ecol. Indic., 60, 213–222, doi:10.1016/j.ecolind.2015.06.043, 2016.

Guisan, A. and Thuiller, W.: Predicting species distribution: offering more than simple habitat models, Ecol. Lett., 8(9), 993–1009, doi:10.1111/j.1461-0248.2005.00792.x, 2005.

Guisan, A. and Zimmermann, N. E.: Predictive habitat distribution models in ecology, Ecol. Model., 135(2-3), 147–186, doi:10.1016/S0304-3800(00)00354-9, 2000.

Harfoot, M., Tittensor, D. P., Newbold, T., McInerny, G., Smith, M. J. and Scharlemann, J. P. W.: Integrated assessment models for ecologists: the present and the future: Integrated assessment models for ecologists, Glob. Ecol. Biogeogr., 23(2), 124–143, doi:10.1111/geb.12100, 2014a.

Harfoot, M. B. J., Newbold, T., Tittensor, D. P., Emmott, S., Hutton, J., Lyutsarev, V., Smith, M. J., Scharlemann, J. P. W. and Purves, D. W.: Emergent Global Patterns of Ecosystem Structure and Function from a Mechanistic General Ecosystem Model, edited by M. Loreau, PLoS Biol., 12(4), e1001841, doi:10.1371/journal.pbio.1001841, 2014b.

Haverd, V., Smith, B., Nieradzik, L., Briggs, P. R., Woodgate, W., Trudinger, C. M. and Canadell, J. G.: A new version of the CABLE land surface model (Subversion revision r4546), incorporating land use and land cover change, woody vegetation demography and a novel optimisation-based approach to plant coordination of electron transport and carboxylation capacity-limited photosynthesis, Geosci. Model Dev. Discuss., 1–33, doi:10.5194/gmd-2017-265, 2017.

Heinimann, A., Mertz, O., Frolking, S., Egelund Christensen, A., Hurni, K., Sedano, F., Parsons Chini, L., Sahajpal, R., Hansen, M. and Hurtt, G.: A global view of shifting cultivation: Recent, current, and future extent, edited by B. Poulter, PLOS ONE, 12(9), e0184479, doi:10.1371/journal.pone.0184479, 2017.

Hempel, S., Frieler, K., Warszawski, L., Schewe, J. and Piontek, F.: A trend-preserving bias correction – the ISI-MIP approach, Earth Syst. Dyn., 4(2), 219–236, doi:10.5194/esd-4-219-2013, 2013.

Hirsch, T. and Secretariat of the Convention on Biological Diversity, Eds.: Global biodiversity outlook 3, Secretariat of the Convention on Biological Diversity, Montreal, Quebec, Canada., 2010.

Hoskins, A. J., Harwood, T.D., Ware, C., Williams, K.J., Perry, J.J., Ota, N., Croft, J.R., Yeates, D.K., Jetz, W., Golebiewski, M., Ferrier, S.: BILBI: supporting global biodiversity assessment through high-resolution macroecological modelling. [in prep.]

Hudson, L. N., Newbold, T., Contu, S., Hill, S. L. L., Lysenko, I., De Palma, A., Phillips, H. R. P., Senior, R. A., Bennett, D. J., Booth, H., Choimes, A., Correia, D. L. P., Day, J., Echeverría-Londoño, S., Garon, M., Harrison, M. L. K., Ingram, D. J., Jung, M., Kemp, V., Kirkpatrick, L., Martin, C. D., Pan, Y., White, H. J., Aben, J., Abrahamczyk, S., Adum, G. B., Aguilar-Barquero, V., Aizen, M. A., Ancrenaz, M., Arbeláez-Cortés, E., Armbrecht, I., Azhar, B., Azpiroz, A. B., Baeten, L., Báldi, A., Banks, J. E., Barlow, J., Batáry, P., Bates, A. J., Bayne, E. M., Beja, P., Berg, Å., Berry, N. J., Bicknell, J. E., Bihn, J. H., Böhning-Gaese, K., Boekhout, T., Boutin, C., Bouyer, J., Brearley, F. Q., Brito, I., Brunet, J., Buczkowski, G., Buscardo, E., Cabra-García, J., Calviño-Cancela, M., Cameron, S. A., Cancello, E. M., Carrijo, T. F., Carvalho, A. L., Castro, H., Castro-Luna, A. A., Cerda, R., Cerezo, A., Chauvat, M., Clarke, F. M., Cleary, D. F. R., Connop, S. P., D’Aniello, B., da Silva, P. G., Darvill, B., Dauber, J., Dejean, A., Diekötter, T., Dominguez-Haydar, Y., Dormann, C. F., Dumont, B., Dures, S. G., Dynesius, M., Edenius, L., Elek, Z., Entling, M. H., Farwig, N., Fayle, T. M., Felicioli, A., Felton, A. M., Ficetola, G. F., Filgueiras, B. K. C., Fonte, S. J., Fraser, L. H., Fukuda, D., Furlani, D., Ganzhorn, J. U., Garden, J. G., Gheler-Costa, C., Giordani, P., Giordano, S., Gottschalk, M.S., Goulson, D., et al.: The PREDICTS database: a global database of how local terrestrial biodiversity responds to human impacts, Ecol. Evol., 4(24), 4701–4735, doi:10.1002/ece3.1303, 2014.

Hurtt, G. C., Chini, L. P., Frolking, S., Betts, R. A., Feddema, J., Fischer, G., Fisk, J. P., Hibbard, K., Houghton, R. A., Janetos, A., Jones, C. D., Kindermann, G., Kinoshita, T., Klein Goldewijk, K., Riahi, K., Shevliakova, E., Smith, S., Stehfest, E., Thomson, A., Thornton, P., van Vuuren, D. P. and Wang, Y. P.: Harmonization of land-use scenarios for the period 1500–2100: 600 years of global gridded annual land-use transitions, wood harvest, and resulting secondary lands, Clim. Change, 109(1–2), 117–161, doi:10.1007/s10584-011-0153-2, 2011.

Hurtt, G., Chini, L., Sahajpal, R., Frolking, S., Calvin, K., Fujimori, S., Klein Goldewijk, K., Hasegawa, T., Havlik, P., Lawrence, D., Lawrence, P., Popp, A., Stehfest, E., van Vuuren, D., and Zhang, X.: Harmonization of global land-use change and management for the period 850–2100. [submitted]

IPBES: The methodological assessment report on scenarios and models of biodiversity and ecosystem services: S. Ferrier, K. N. Ninan, P. Leadley, R. Alkemade, L. A. Acosta, H. R. Akçakaya, L. Brotons, W. W. L. Cheung, V. Christensen, K. A. Harhash, J. Kabubo-Mariara, C. Lundquist, M. Obersteiner, H. M. Pereira, G. Peterson, R. Pichs-Madruga, N. Ravindranath, C. Rondinini and B. A. Wintle (eds.), Secretariat of the Intergovernmental Science-Policy Platform on Biodiversity and Ecosystem Services, Bonn, Germany, 348 pages, 2016.

Janse J.H., M. Bakkenes & J. Meijer: Globio-Aquatic, Technical model description v. 1.3: PBL publication 2829, The Hague, PBL Netherlands Environmental Assessment Agency, 2016.

Janse, J. H., Kuiper, J. J., Weijters, M. J., Westerbeek, E. P., Jeuken, M. H. J. L., Bakkenes, M., Alkemade, R., Mooij, W. M. and Verhoeven, J. T. A.: GLOBIO-Aquatic, a global model of human impact on the biodiversity of inland aquatic ecosystems, Environ. Sci. Policy, 48, 99–114, doi:10.1016/j.envsci.2014.12.007, 2015.

Jantz, S. M., Barker, B., Brooks, T. M., Chini, L. P., Huang, Q., Moore, R. M., Noel, J. and Hurtt, G. C.: Future habitat loss and extinctions driven by land-use change in biodiversity hotspots under four scenarios of climate-change mitigation: Future Habitat Loss and Extinctions, Conserv. Biol., 29(4), 1122–1131, doi:10.1111/cobi.12549, 2015.

Jetz, W., Wilcove, D. S. and Dobson, A. P.: Projected Impacts of Climate and Land-Use Change on the Global Diversity of Birds, edited by G. M. Mace, PLoS Biol., 5(6), e157, doi:10.1371/journal.pbio.0050157, 2007.

Johnson, J. A., Runge, C. F., Senauer, B., Foley, J. and Polasky, S.: Global agriculture and carbon trade-offs, Proc. Natl. Acad. Sci., 111(34), 12342–12347, doi:10.1073/pnas.1412835111, 2014.

Johnson, J. A., Runge, C. F., Senauer, B. and Polasky, S.: Global Food Demand and Carbon-Preserving Cropland Expansion under Varying Levels of Intensification, Land Econ., 92(4), 579–592, doi:10.3368/le.92.4.579, 2016.

Jungclaus, J. H., Bard, E., Baroni, M., Braconnot, P., Cao, J., Chini, L. P., Egorova, T., Evans, M., González-Rouco, J. F., Goosse, H., Hurtt, G. C., Joos, F., Kaplan, J. O., Khodri, M., Klein Goldewijk, K., Krivova, N., LeGrande, A. N., Lorenz, S. J., Luterbacher, J., Man, W., Maycock, A. C., Meinshausen, M., Moberg, A., Muscheler, R., Nehrbass-Ahles, C., Otto-Bliesner, B. I., Phipps, S. J., Pongratz, J., Rozanov, E., Schmidt, G. A., Schmidt, H., Schmutz, W., Schurer, A., Shapiro, A. I., Sigl, M., Smerdon, J. E., Solanki, S. K., Timmreck, C., Toohey, M., Usoskin, I. G., Wagner, S., Wu, C.-J., Yeo, K. L., Zanchettin, D., Zhang, Q. and Zorita, E.: The PMIP4 contribution to CMIP6 – Part 3: The last millennium, scientific objective, and experimental design for the PMIP4 *past1000* simulations, Geosci. Model Dev., 10(11), 4005–4033, doi:10.5194/gmd-10-4005-2017, 2017.

Knorr, W., Arneth, A. and Jiang, L.: Demographic controls of future global fire risk, Nat. Clim. Change, 6(8), 781 – 785, doi:10.1038/nclimate2999, 2016.

Kriegler, E., Bauer, N., Popp, A., Humpenöder, F., Leimbach, M., Strefler, J., Baumstark, L., Bodirsky, B. L., Hilaire, J., Klein, D., Mouratiadou, I., Weindl, I., Bertram, C., Dietrich, J.-P., Luderer, G., Pehl, M., Pietzcker, R., Piontek, F., Lotze-Campen, H., Biewald, A., Bonsch, M., Giannousakis, A., Kreidenweis, U., Müller, C., Rolinski, S., Schultes, A., Schwanitz, J., Stevanovic, M., Calvin, K., Emmerling, J., Fujimori, S. and Edenhofer, O.: Fossil-fueled development (SSP5): An energy and resource intensive scenario for the 21st century, Glob. Environ. Change, 42, 297–315, doi:10.1016/j.gloenvcha.2016.05.015, 2017.

Lawrence, D. M., Hurtt, G. C., Arneth, A., Brovkin, V., Calvin, K. V., Jones, A. D., Jones, C. D., Lawrence, P. J., de Noblet-Ducoudré, N., Pongratz, J., Seneviratne, S. I. and Shevliakova, E.: The Land Use Model Intercomparison Project (LUMIP) contribution to CMIP6: rationale and experimental design, Geosci. Model Dev., 9(9), 2973–2998, doi:10.5194/gmd-9-2973-2016, 2016.

Leadley, P. W., Krug, C.B., Alkemade, R., Pereira, H.M., Sumaila U.R., Walpole, M., Marques, A., Newbold, T., Teh, L.S.L, van Kolck, J., Bellard, C., Januchowski-Hartley, S.R. and Mumby, P.J.: Progress towards the Aichi Biodiversity Targets: An Assessment of Biodiversity Trends, Policy Scenarios and Key Actions. Secretariat of the Convention on Biological Diversity, Montreal, Canada, Technical Series 78, 500 pages, 2014.

Lehsten, V., Sykes, M. T., Scott, A. V., Tzanopoulos, J., Kallimanis, A., Mazaris, A., Verburg, P. H., Schulp, C. J. E., Potts, S. G. and Vogiatzakis, I.: Disentangling the effects of land-use change, climate and CO 2 on projected future European habitat types: Disentangling the drivers of habitat change, Glob. Ecol. Biogeogr., 24(6), 653–663, doi:10.1111/geb.12291, 2015.

Lindeskog, M., Ameth, A., Bondeau, A., Waha, K., Seaquist, J., Olin, S. and Smith, B.: Implications of accounting for land use in simulations of ecosystem carbon cycling in Africa, Earth Syst. Dyn., 4(2), 385–407, doi:10.5194/esd-4-385-2013, 2013.

Martins, I. S. and Pereira, H. M.: Improving extinction projections across scales and habitats using the countryside species-area relationship, Sci. Rep., 7(1), doi:10.1038/s41598-017-13059-y, 2017.

Maxwell, S. L., Fuller, R. A., Brooks, T. M. and Watson, J. E. M.: Biodiversity: The ravages of guns. nets and bulldozers, Nature, 536(7615), 143–145, doi:10.1038/536143a, 2016.

McSweeney, C. F. and Jones, R. G.: How representative is the spread of climate projections from the 5 CMIP5 GCMs used in ISI-MIP?, Clim. Serv., 1, 24–29, doi:10.1016/j.cliser.2016.02.001, 2016.

Meinshausen, M., Wigley, T. M. L. and Raper, S. C. B.: Emulating atmosphere-ocean and carbon cycle models with a simpler model, MAGICC6 – Part 2: Applications, Atmospheric Chem. Phys., 11(4), 1457–1471, doi:10.5194/acp-11-1457-2011, 2011a.

Meinshausen, M., Raper, S. C. B. and Wigley, T. M. L.: Emulating coupled atmosphere-ocean and carbon cycle models with a simpler model, MAGICC6 – Part 1: Model description and calibration, Atmospheric Chem. Phys., 11(4), 1417–1456, doi:10.5194/acp-11-1417-2011, 2011b.

Merow, C., Smith, M. J. and Silander, J. A.: A practical guide to MaxEnt for modeling species’ distributions: what it does, and why inputs and settings matter, Ecography, 36(10), 1058–1069, doi:10.1111/j.1600-0587.2013.07872.x, 2013.

Millennium Ecosystem Assessment (Program), Ed.: Ecosystems and human well-being: synthesis, Island Press, Washington, DC., 2005.

Monfreda, C., Ramankutty, N. and Foley, J. A.: Farming the planet: 2. Geographic distribution of crop areas, yields, physiological types, and net primary production in the year 2000: GLOBAL CROP AREAS AND YIELDS IN 2000, Glob. Biogeochem. Cycles, 22(1), n/a–n/a, doi:10.1029/2007GB002947, 2008.

Moss, R. H., Edmonds, J. A., Hibbard, K. A., Manning, M. R., Rose, S. K., van Vuuren, D. P., Carter, T. R., Emori, S., Kainuma, M., Kram, T., Meehl, G. A., Mitchell, J. F. B., Nakicenovic, N., Riahi, K., Smith, S. J., Stouffer, R. J., Thomson, A. M., Weyant, J. P. and Wilbanks, T. J.: The next generation of scenarios for climate change research and assessment, Nature, 463(7282), 747–756, doi:10.1038/nature08823, 2010.

Newbold, T., Hudson, L. N., Arnell, A. P., Contu, S., De Palma, A., Ferrier, S., Hill, S. L. L., Hoskins, A. J., Lysenko, I., Phillips, H. R. P., Burton, V. J., Chng, C. W. T., Emerson, S., Gao, D., Pask-Hale, G., Hutton, J., Jung, M., Sanchez-Ortiz, K., Simmons, B. I., Whitmee, S., Zhang, H., Scharlemann, J. P. W. and Purvis, A.: Has land use pushed terrestrial biodiversity beyond the planetary boundary? A global assessment, Science, 353(6296), 288–291, doi:10.1126/science.aaf2201, 2016.

Ohashi, H., Hasegawa, T., Hirata, A., Fujimori, S., Takahashi, K., Hijioka, Y., Nakao, K., Tsuyama, I., Kominami, Y., Tanaka, N., Matsui, T.: Long-term climate change impact is more harmful than 2°C climate mitigation land use change. [in prep.]

Olin, S., Schurgers, G., Lindeskog, M., Wårlind, D., Smith, B., Bodin, P., Holmér, J. and Arneth, A.: Modelling the response of yields and tissue C : N to changes in atmospheric CO_2_ and N management in the main wheat regions of western Europe, Biogeosciences, 12(8), 2489–2515, doi:10.5194/bg-12-2489-2015, 2015.

O’Neill, B. C., Tebaldi, C., van Vuuren, D. P., Eyring, V., Friedlingstein, P., Hurtt, G., Knutti, R., Kriegler, E., Lamarque, J.-F., Lowe, J., Meehl, G. A., Moss, R., Riahi, K. and Sanderson, B. M.: The Scenario Model Intercomparison Project (ScenarioMIP) for CMIP6, Geosci. Model Dev., 9(9), 3461–3482, doi:10.5194/gmd-9-3461-2016, 2016.

O’Neill, B. C., Kriegler, E., Ebi, K. L., Kemp-Benedict, E., Riahi, K., Rothman, D. S., van Ruijven, B. J., van Vuuren, D. P., Birkmann, J., Kok, K., Levy, M. and Solecki, W.: The roads ahead: Narratives for shared socioeconomic pathways describing world futures in the 21st century, Glob. Environ. Change, 42, 169–180, doi:10.1016/j.gloenvcha.2015.01.004, 2017.

Pecl, G. T., Araújo, M. B., Bell, J. D., Blanchard, J., Bonebrake, T. C., Chen, I.-C., Clark, T. D., Colwell, R. K., Danielsen, F., Evengård, B., Falconi, L., Ferrier, S., Frusher, S., Garcia, R. A., Griffis, R. B., Hobday, A. J., Janion-Scheepers, C., Jarzyna, M. A., Jennings, S., Lenoir, J., Linnetved, H. I., Martin, V. Y., McCormack, P. C., McDonald, J., Mitchell, N. J., Mustonen, T., Pandolfi, J. M., Pettorelli, N., Popova, E., Robinson, S. A., Scheffers, B. R., Shaw, J. D., Sorte, C. J. B., Strugnell, J. M., Sunday, J. M., Tuanmu, M.-N., Vergés, A., Villanueva, C., Wernberg, T., Wapstra, E. and Williams, S. E.: Biodiversity redistribution under climate change: Impacts on ecosystems and human well-being, Science, 355(6332), eaai9214, doi:10.1126/science.aai9214, 2017.

Pereira, H. M., Leadley, P. W., Proenca, V., Alkemade, R., Scharlemann, J. P. W., Fernandez-Manjarres, J. F., Araujo, M. B., Balvanera, P., Biggs, R., Cheung, W. W. L., Chini, L., Cooper, H. D., Gilman, E. L., Guenette, S., Hurtt, G. C., Huntington, H. P., Mace, G. M., Oberdorff, T., Revenga, C., Rodrigues, P., Scholes, R. J., Sumaila, U. R. and Walpole, M.: Scenarios for Global Biodiversity in the 21st Century, Science, 330(6010), 1496–1501, doi:10.1126/science.1196624, 2010.

Popp, A., Calvin, K., Fujimori, S., Havlik, P., Humpenöder, F., Stehfest, E., Bodirsky, B. L., Dietrich, J. P., Doelmann, J. C., Gusti, M., Hasegawa, T., Kyle, P., Obersteiner, M., Tabeau, A., Takahashi, K., Valin, H., Waldhoff, S., Weindl, I., Wise, M., Kriegler, E., Lotze-Campen, H., Fricko, O., Riahi, K. and Vuuren, D. P. van: Land-use futures in the shared socio-economic pathways, Glob. Environ. Change, 42, 331–345, doi:10.1016/j.gloenvcha.2016.10.002, 2017.

Poulter, B., Frank, D. C., Hodson, E. L. and Zimmermann, N. E.: Impacts of land cover and climate data selection on understanding terrestrial carbon dynamics and the CO_2_ airborne fraction, Biogeosciences, 8(8), 2027–2036, doi:10.5194/bg-8-2027-2011, 2011.

Prentice, I. C., Bondeau, A., Cramer, W., Harrison, S. P., Hickler, T., Lucht, W., Sitch, S., Smith, B. and Sykes, M. T.: Dynamic Global Vegetation Modeling: Quantifying Terrestrial Ecosystem Responses to Large-Scale Environmental Change, in Terrestrial Ecosystems in a Changing World, D. E. Pataki, and L. F. Pitelka, pp. 175–192, Springer Berlin Heidelberg, Berlin, Heidelberg., 2007.

Purvis, A., Newbold, T., De Palma, A., Contu, S., Hill, S. L. L., Sanchez-Ortiz, K., Phillips, H. R. P., Hudson, L. N., Lysenko, I., Börger, L. and Scharlemann, J. P. W.: Modelling and Projecting the Response of Local Terrestrial Biodiversity Worldwide to Land Use and Related Pressures: The PREDICTS Project, in Advances in Ecological Research, vol. 58, pp. 201–241, Elsevier., 2018.

Rabin, S. S., Melton, J. R., Lasslop, G., Bachelet, D., Forrest, M., Hantson, S., Kaplan, J. O., Li, F., Mangeon, S., Ward, D. S., Yue, C., Arora, V. K., Hickler, T., Kloster, S., Knorr, W., Nieradzik, L., Spessa, A., Folberth, G. A., Sheehan, T., Voulgarakis, A., Kelley, D. I., Prentice, I. C., Sitch, S., Harrison, S. and Arneth, A.: The Fire Modeling Intercomparison Project (FireMIP), phase 1: experimental and analytical protocols with detailed model descriptions, Geosci. Model Dev., 10(3), 1175–1197, doi:10.5194/gmd-10-1175-2017, 2017.

Redhead, J. W., May, L., Oliver, T. H., Hamel, P., Sharp, R. and Bullock, J. M.: National scale evaluation of the InVEST nutrient retention model in the United Kingdom, Sci. Total Environ., 610–611, 666–677, doi:10.1016/j.scitotenv.2017.08.092, 2018.

Riahi, K., van Vuuren, D. P., Kriegler, E., Edmonds, J., O’Neill, B. C., Fujimori, S., Bauer, N., Calvin, K., Dellink, R., Fricko, O., Lutz, W., Popp, A., Cuaresma, J. C., Kc, S., Leimbach, M., Jiang, L., Kram, T., Rao, S., Emmerling, J., Ebi, K., Hasegawa, T., Havlik, P., Humpenöder, F., Da Silva, L. A., Smith, S., Stehfest, E., Bosetti, V., Eom, J., Gernaat, D., Masui, T., Rogelj, J., Strefler, J., Drouet, L., Krey, V., Luderer, G., Harmsen, M., Takahashi, K., Baumstark, L., Doelman, J. C., Kainuma, M., Klimont, Z., Marangoni, G., Lotze-Campen, H., Obersteiner, M., Tabeau, A. and Tavoni, M.: The Shared Socioeconomic Pathways and their energy, land use, and greenhouse gas emissions implications: An overview, Glob. Environ. Change, 42, 153–168, doi:10.1016/j.gloenvcha.2016.05.009, 2017.

Rondinini, C., Di Marco, M., Chiozza, F., Santulli, G., Baisero, D., Visconti, P., Hoffmann, M., Schipper, J., Stuart, S. N., Tognelli, M. F., Amori, G., Falcucci, A., Maiorano, L. and Boitani, L.: Global habitat suitability models of terrestrial mammals, Philos. Trans. R. Soc. B Biol. Sci., 366(1578), 2633–2641, doi:10.1098/rstb.2011.0113, 2011.

Rosa, I. M., Pereira, H. M., Ferrier, S., Alkemade, R., Acosta, L. A., Akcakaya, H. R., den Belder, E., Fazel, A. M., Fujimori, S. and Harfoot, M.: Multiscale scenarios for nature futures, Nat. Ecol. Evol., 1(10), 1416, 2017.

Rosenzweig, C., Arnell, N. W., Ebi, K. L., Lotze-Campen, H., Raes, F., Rapley, C., Smith, M. S., Cramer, W., Frieler, K., Reyer, C. P. O., Schewe, J., van Vuuren, D. and Warszawski, L.: Assessing inter-sectoral climate change risks: the role of ISIMIP, Environ. Res. Lett., 12(1), 010301, doi:10.1088/1748-9326/12/1/010301, 2017.

Sala, O. E.: Global Biodiversity Scenarios for the Year 2100&nbsp;, Science, 287(5459), 1770–1774, doi:10.1126/science.287.5459.1770, 2000.

Schipper, A. M., Bakkenes, M., Meijer, J.R., Alkemade, R., Huijbregts, M.J.: The GLOBIO model. A technical description of version 3.5. PBL publication 2369, The Hague, PBL Netherlands Environmental Assessment Agency, 2016.

Schulp, C. J. E., Alkemade, R., Klein Goldewijk, K. and Petz, K.: Mapping ecosystem functions and services in Eastern Europe using global-scale data sets, Int. J. Biodivers. Sci. Ecosyst. Serv. Manag., 8(1–2), 156–168, doi:10.1080/21513732.2011.645880, 2012.

Secretariat of the Convention on Biological Diversity and United Nations Environment Programme, Eds.: Global biodiversity outlook 4: a mid-term assessment of progress towards the implementation of the strategic plan for biodiversity 2011-2020, Secretariat for the Convention on Biological Diversity, Montreal, Quebec, Canada., 2014.

Settele, J., Scholes, R., Betts, R. A., Bunn, S., Leadley, P., Nepstad, D., … Winter, M.: Terrestrial and Inland water systems: in, Climate Change 2014 Impacts, Adaptation and Vulnerability: Part A: Global and Sectoral Aspects (pp. 271–360), Cambridge University Press, DOI: 10.1017/CBO9781107415379.009, 2015.

Sharp, R., Tallis, H.T., Ricketts, T., Guerry, A.D., Wood, S.A., Chaplin-Kramer, R., Nelson, E., Ennaanay, D., Wolny, S., Olwero, N., Vigerstol, K., Pennington, D., Mendoza, G., Aukema, J., Foster, J., Forrest, J., Cameron, D., Arkema, K., Lonsdorf, E., Kennedy, C., Verutes, G., Kim, C.K., Guannel, G., Papenfus, M., Toft, J., Marsik, M., Bernhardt, J., Griffin, R., Glowinski, K., Chaumont, N., Perelman, A., Lacayo, M. Mandle, L., Hamel, P., Vogl, A.L., Rogers, L., Bierbower, W., Denu, D., and Douglass, J.: InVEST +VERSION+ User’s Guide, The Natural Capital Project, Stanford University, University of Minnesota, The Nature Conservancy, and World Wildlife Fund, 2018.

Sitch, S., Smith, B., Prentice, I. C., Arneth, A., Bondeau, A., Cramer, W., Kaplan, J. O., Levis, S., Lucht, W., Sykes, M. T., Thonicke, K. and Venevsky, S.: Evaluation of ecosystem dynamics, plant geography and terrestrial carbon cycling in the LPJ dynamic global vegetation model, Glob. Change Biol., 9(2), 161–185, doi:10.1046/j.1365-2486.2003.00569.x, 2003.

Smith, B., Wårlind, D., Arneth, A., Hickler, T., Leadley, P., Siltberg, J. and Zaehle, S.: Implications of incorporating N cycling and N limitations on primary production in an individual-based dynamic vegetation model, Biogeosciences, 11(7), 2027–2054, doi:10.5194/bg-11-2027-2014, 2014.

Stehfest, E., van Vuuren, D., Kram, T., Bouwman, L., Alkemade, R., Bakkenes, M., Biemans, H., Bouwman, A., den Elzen, M., Janse, J., Lucas, P., van Minnen, J., Müller, M., Prins, A.: Integrated Assessment of Global Environmental Change with IMAGE 3.0. Model description and policy applications, The Hague: PBL Netherlands Environmental Assessment Agency, 2014.

Thuiller, W.: Patterns and uncertainties of species’ range shifts under climate change, Glob. Change Biol., 10(12), 2020–2027, doi:10.1111/j.1365-2486.2004.00859.x, 2004.

Thuiller, W., Lafourcade, B., Engler, R. and Araújo, M. B.: BIOMOD - a platform for ensemble forecasting of species distributions, Ecography, 32(3), 369–373, doi:10.1111/j.1600-0587.2008.05742.x, 2009.

Thuiller, W., Lavergne, S., Roquet, C., Boulangeat, I., Lafourcade, B. and Araujo, M. B.: Consequences of climate change on the tree of life in Europe, Nature, 470(7335), 531–534, doi:10.1038/nature09705, 2011.

Thuiller, W., Münkemüller, T., Lavergne, S., Mouillot, D., Mouquet, N., Schiffers, K. and Gravel, D.: A road map for integrating eco-evolutionary processes into biodiversity models, edited by M. Holyoak, Ecol. Lett., 16, 94–105, doi:10.1111/ele.12104, 2013.

Titeux, N., Henle, K., Mihoub, J.-B., Regos, A., Geijzendorffer, I. R., Cramer, W., Verburg, P. H. and Brotons, L.: Biodiversity scenarios neglect future land-use changes, Glob. Change Biol., 22(7), 2505–2515, doi:10.1111/gcb.13272, 2016.

Titeux, N., Henle, K., Mihoub, J.-B., Regos, A., Geijzendorffer, I. R., Cramer, W., Verburg, P. H. and Brotons, L.: Global scenarios for biodiversity need to better integrate climate and land use change, edited by A. Syphard, Divers. Distrib., 23(11), 1231–1234, doi:10.1111/ddi.12624, 2017.

United Nations Environment Programme: UNEP-SETAC Life Cycle Initiative: Global Guidance for Life Cycle Impact Assessment Indicators - Volume 1, 2016.

Visconti, P., Bakkenes, M., Baisero, D., Brooks, T., Butchart, S. H. M., Joppa, L., Alkemade, R., Di Marco, M., Santini, L., Hoffmann, M., Maiorano, L., Pressey, R. L., Arponen, A., Boitani, L., Reside, A. E., van Vuuren, D. P. and Rondinini, C.: Projecting Global Biodiversity Indicators under Future Development Scenarios: Projecting biodiversity indicators, Conserv. Lett., 9(1), 5–13, doi:10.1111/conl.12159, 2016.

van Vuuren, D. P., Edmonds, J., Kainuma, M., Riahi, K., Thomson, A., Hibbard, K., Hurtt, G. C., Kram, T., Krey, V., Lamarque, J.-F., Masui, T., Meinshausen, M., Nakicenovic, N., Smith, S. J. and Rose, S. K.: The representative concentration pathways: an overview, Clim. Change, 109(1–2), 5–31, doi:10.1007/s10584-011-0148-z, 2011.

van Vuuren, D. P., Kriegler, E., O’Neill, B. C., Ebi, K. L., Riahi, K., Carter, T. R., Edmonds, J., Hallegatte, S., Kram, T., Mathur, R. and Winkler, H.: A new scenario framework for Climate Change Research: scenario matrix architecture, Clim. Change, 122(3), 373–386, doi:10.1007/s10584-013-0906-1, 2014.

van Vuuren, D. P., Stehfest, E., Gernaat, D. E. H. J., Doelman, J. C., van den Berg, M., Harmsen, M., de Boer, H. S., Bouwman, L. F., Daioglou, V., Edelenbosch, O. Y., Girod, B., Kram, T., Lassaletta, L., Lucas, P. L., van Meijl, H., Müller, C., van Ruijven, B. J., van der Sluis, S. and Tabeau, A.: Energy, land-use and greenhouse gas emissions trajectories under a green growth paradigm, Glob. Environ. Change, 42, 237–250, doi:10.1016/j.gloenvcha.2016.05.008, 2017.

Warszawski, L., Frieler, K., Huber, V., Piontek, F., Serdeczny, O. and Schewe, J.: The Inter-Sectoral Impact Model Intercomparison Project (ISI–MIP): Project framework, Proc. Natl. Acad. Sci., 111(9), 3228–3232, doi:10.1073/pnas.1312330110, 2014.

Zhao, J.: Influencing policymakers: Communication, Nat. Clim. Change, 7(3), 173–174, doi:10.1038/nclimate3215, 2017.

